# A flexible workflow for simulating transcranial electric stimulation in healthy and lesioned brains

**DOI:** 10.1101/2020.01.09.900035

**Authors:** Benjamin Kalloch, Pierre-Louis Bazin, Arno Villringer, Bernhard Sehm, Mario Hlawitschka

## Abstract

Simulating transcranial electric stimulation is actively researched as knowledge about the distribution of the electrical field is decisive for understanding the variability in the elicited stimulation effect. Several software pipelines comprehensively solve this task in an automated manner for standard use-cases. However, simulations for non-standard applications such as uncommon electrode shapes or the creation of head models from non-optimized T1-weighted imaging data and the inclusion of irregular structures are more difficult to accomplish.

We address these limitations and suggest a comprehensive workflow to simulate transcranial electric stimulation based on open-source tools. The workflow covers the head model creation from MRI data, the electrode modeling, the modeling of anisotropic conductivity behavior of the white matter, the numerical simulation and visualization.

Skin, skull, air cavities, cerebrospinal fluid, white matter, and gray matter are segmented semi-automatically from T1-weighted MR images. Electrodes of arbitrary number and shape can be modeled. The meshing of the head model is implemented in a way to preserve feature edges of the electrodes and is free of topological restrictions of the considered structures of the head model. White matter anisotropy can be computed from diffusion-tensor imaging data.

Our solver application was verified analytically and by contrasting tDCS simulation results with another simulation pipeline (SimNIBS 3.0). An agreement in both cases underlines the validity of our workflow.

Our suggested solutions facilitate investigations of irregular structures in patients (e.g. lesions, implants) or of new electrode types. For a coupled use of the described workflow, we provide documentation and disclose the full source code of the developed tools.

## 1. Introduction

The simulation of transcranial electric stimulation (tES) is increasingly employed when designing tES intervention studies (1) and observed behavior or neurophysiological changes are related to the simulated, subject-specific electric field (2), (3), (4). This development is motivated by increasing evidence that the individual distribution of the electrical field within each subject influences the stimulation effect (5), (6), (7). In addition, several software pipelines (8), (9), (10), (11), (12), among which SimNIBS (10) and ROAST (12) are currently most actively developed, make the simulation of tES more accessible to researchers.

All these pipelines implement a common, general workflow covering standard use cases, i.e. the tES simulation of healthy subjects based on their individual magnetic resonance imaging (MRI) data using rectangular or circular electrodes. The starting point of this workflow is the segmentation of the MRI data of the subjects into the electrically most important tissue classes. The obtained segmentation image is then used to create the head volume mesh, which is complemented by electrodes that need to be modeled and positioned. The simulation problem is solved using this individual head model, and results are visualized. The implementation of the outlined workflow by current tES simulation pipelines does not entirely cover use cases with suboptimal imaging data, the presence of pathological tissue in patients or alternative electrode shapes.

For instance, MRI data from large-scale imaging studies usually have not been primarily acquired for the purpose of computational head modeling. Performing simulation studies based on such data can become difficult due to challenges in the segmentation of low-contrast tissue such as skull using standard segmentation approaches. Following the image segmentation, a surface-based meshing approach is commonly used to create the head volume mesh. The advantage of this approach is a maximum of control over the approximation of the boundaries of the sub-compartments of the head model, which, on the other hand, must not intersect, restricting the topology of the included structures and complicating the inclusion of irregular tissue such as lesioned tissue. ROAST circumvents this restriction by applying an image-based meshing approach, which is free of any topological constraints (12), with the drawback of less accurate feature edges, for example of the electrodes. The shape of the electrodes commonly can be selected from a set of standard shapes including rectangular, circular or ring electrodes. Means for modeling non-standard shaped electrodes such triangular electrodes are usually not provided. Finally, the visualization of the simulation results is typically realized in MATLAB (8), (12), (11), GMSH (13) or a custom tool (9) and thus relatively limited.

In this work, we present approaches to address the above-mentioned non-standard use-cases when simulating tES on an individual basis. Segmentation routines were selected based on the robustness of the structure segmentation of T1-weighted MRI data using JIST (14) a plugin of MIPAV (15) to benefit from a wide range of image manipulation and segmentation algorithms. We introduce an extension to the image-based meshing approach presented in (12) by combining it with a surface-based meshing approach for an accurate electrode representation. The 3D modeling software Blender (16) allows highly flexible modeling of electrodes of arbitrary shapes. We suggest the use of ParaView (17) for a versatile visualization of the simulation results. We describe the information flow among the involved tools, which are arranged around OpenFOAM (18), a comprehensive, finite-volume-method-based framework for the numerical simulations. The simulation was verified analytically and by contrasting the numerical results with those of SimNIBS 3.0. A general agreement between both approaches underlines the validity of our suggested solutions. The scripts and the custom source code along with the documentation are readily available (from https://gitlab.imn.htwk-leipzig.de/bkalloc1/tdcs-pipeline.git) allowing a coupled used of the entire toolset as well as usage of single tools only.

## 2. Methods

The process of simulating tES involves the head and the electrode modeling, solving the underlying electrostatic problem, and the visualization.

The head model creation comprises the segmentation of the head MR image and the subsequent volume mesh generation. Here, image segmentation is performed using the Java Image Science Toolkit (JIST) (14) a plugin of the Medical Image Processing, Analysis, and Visualization (MIPAV) toolbox (15). The volume mesh is generated using a combined image- and surface-based meshing approach implemented as a custom application that uses the *Computational Geometry Algorithms Library* (CGAL) API, version 4.13.1 (19). A plugin for the 3D modeling software Blender 2.79 (16) implements the modeling and positioning of the electrodes. OpenFOAM 7.0 (18) provides the tools to define the conductivity values of the mesh compartments. Additionally, information from diffusion-weighted imaging (DWI) data can be incorporated to model the anisotropic conducting behavior of white matter tissue, and are processed in MRTrix 3 (20). A plugin developed for the visualization software ParaView 5.6 (21) manages the calculation of the conductivity tensors derived from the diffusion tensors. The finite volume calculations involved in solving the underlying Maxwell’s equation are performed by a custom solver application implementing the OpenFOAM API. Finally, the resulting electric field may be visualized in ParaView. Fig. 1 illustrates the entire workflow.

**Fig 1.**
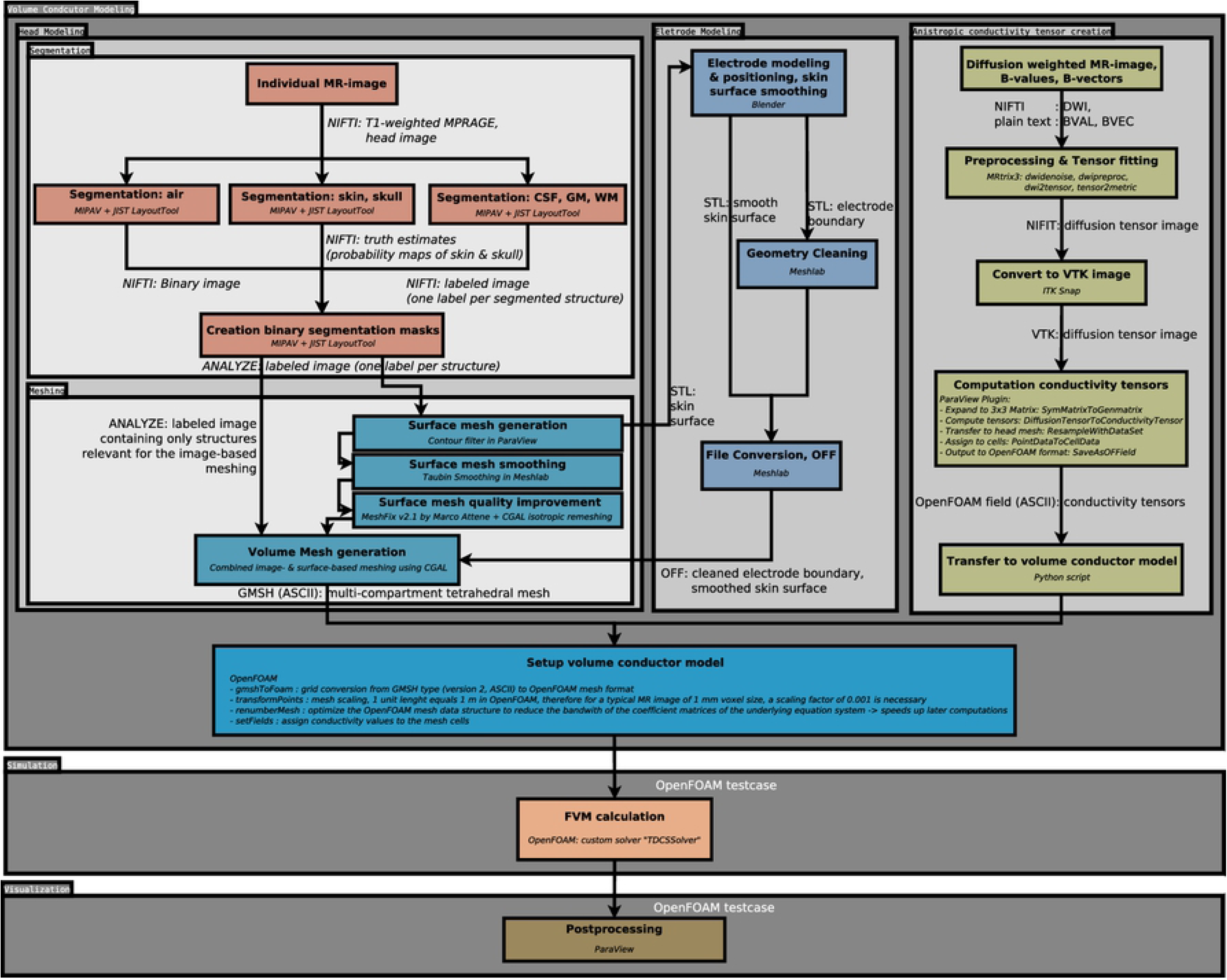
Data- and workflow. Schematics depicting the dataflow between the individual processing steps and the involved tools as well as the expected input and output data of the individual stages.

### 2.1 Set-up of the volume conductor model

#### 2.1.1 MRI head segmentation

Accurate segmentation of the MR image is crucial since the segmented structures represent the individual compartments of the volume conductor model. Segmentation errors - especially discontinuities of the segmented skull or cerebrospinal fluid (CSF) - impair the simulation results (22). In our approach, we segment the scalp, the skull, the air-filled sinuses of the skull, the subarachnoid CSF, the CSF in the ventricles, the gray matter (GM) and the white matter (WM) only from T1-weighted MRI data. The involved segmentation process is described in our previous work (23). In short, we rely on robust, atlas-based segmentation techniques and image-processing capabilities implemented in JIST, a plugin of MIPAV. The segmentation of the scalp and skull structure of the image is achieved through the *Simultaneous Truth And Performance Level Estimation* algorithm (24). The intracranial compartments are segmented using the topology-preserving segmentation algorithm *Multi-object Geometric Deformable Model* (25) and the gyrification of the segmented GM surface is enhanced by the *Cortical Reconstruction Using Implicit Surface Evolution* method (26). We use a pseudo-CT template (27) to segment the air cavities in the skull. The quality of the generated segmentation images is improved by morphological image operations. The individual segmentation images are combined to a single image that contains a distinct, unique numeric label per segmented structure and is exported in the ANALYZE file format.

#### 2.1.2 Electrode modeling and positioning

In our workflow a complete electrode model is implemented, which defines the electrodes geometrically in shape and position as well as their physical conductivity and the applied current, thereby realistically modeling the current shunt (28). The power source is represented by equipotential surfaces at the outer boundaries of the electrode. An optional gel layer may be modeled.

A custom Blender plugin geometrically models rectangular electrodes and positions them according to the international 10-20 system in a semi-automatic way. Necessary inputs are 1) a geometrical representation of the outer boundary of the scalp segmentation in the Stereolithography (STL) file format, 2) the extents of the electrode and 3) its location in 10-20 coordinates. Furthermore, the user must provide four fiducial points, namely the nasion, inion and the tragi of the ears, on the scalp surface by interactively aligning two reference lines and selecting the corresponding points on these lines. The user interface is shown in Fig. 2 A.

**Fig 2.**
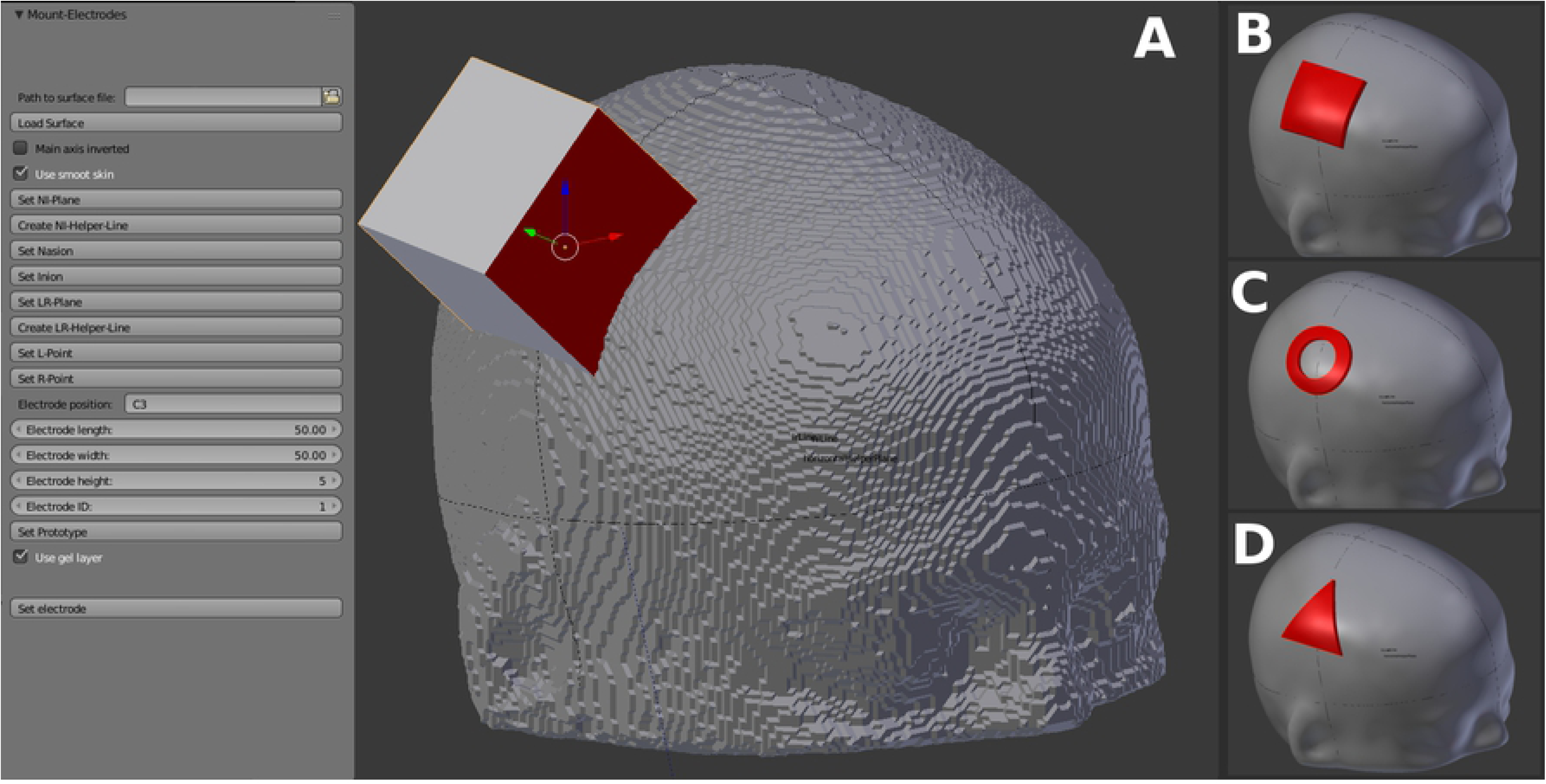
Electrode Modeling. **A)** The user interface of our Blender plugin for electrode positioning and modeling purposes. Necessary input parameters constitute the electrode dimensions, position according to the 10-20 system, and a geometrical representation of the outer boundary of the scalp segmentation in the STL file format. A stepwise workflow to define fiducial points (nasion, inion, tragi of the ears) for the computation of the 10-20 coordinate grid on the individual head is provided by the GUI. The rectangular cube is generated according to the defined dimension and position of the electrode and will be used to create the electrode by the means of constructive solid geometry (CSG). **B)** Result obtained with our plugin: a standard rectangular patch electrode located at C3. A smooth representation of the skin is generated, and the electrode is extruded based on the result of the CSG operation of the cube and this skin surface. **C)** A ring electrode shape created by a non-standard workflow. The cube was replaced by a cylinder with a hole. **D)** Triangular electrode obtained by a non-standard workflow. The cube was replaced by a triangular prism.

To create the geometrical surface representation of the outer scalp boundary from the binary scalp segmentation image the Marching Cubes-based (MC) “Contour Filter” in ParaView is used. In Blender, the plugin initially performs a Laplacian smoothing of the input scalp surface to mitigate its relatively coarse structure due to the MC algorithm. Then, the location of the 10-20 coordinates on the smoothed scalp surface is computed using the user-defined fiducial points. The smooth scalp surface is clipped by the means of constructive solid geometry (CSG) at the specified location with a cube of the specified extent. The position of this cube may be manually varied if the location of the electrode falls outside the standard 10/20 grid. An arbitrary shape of the electrode (see Figs. 2 B - D) can be achieved by replacing that cube with a volume of the desired shape. The clipped surface patch is extruded in 1 mm steps to the desired electrode thickness. This avoids long, thin triangles at the sidewalls of the electrode representation which are unfavorable for the subsequent volume meshing. To model a gel layer this process is executed twice, and the electrode representation is moved on top of the gel layer. The geometry of the electrodes, the gel layer, and the smooth skin surface are exported as STL files.

The CSG operation may result in small, unfavorably clipped triangles at the edges of the electrode and the gel layer that impede the subsequent volume mesh generation. Therefore, their geometry must be cleaned in *Meshlab* (29) by unifying duplicate vertices and applying the “*Quadratic Edge Collapse Decimation*” simplification filter. The smoothed skin surface, the cleaned electrodes, and the gel layer are converted to the Object File Format (OFF).

#### 2.1.3 Volume meshing

An unstructured tetrahedral mesh constitutes the computational domain, i.e. the head model. We approach the task of generating this mesh by applying a combination of an image-based meshing and a surface-based meshing algorithm, both relying on Delaunay triangulation that is implemented in the *Computational Geometry Algorithms Library* (CGAL), version 4.13.1. The surface-based meshing is applied to the electrodes and the scalp structure and can be further utilized for any following internal structure that does not impede a strictly nested arrangement of the mesh compartments. Structures that violate a nested arrangement, such as the ventricles or lesioned tissue, can be meshed using the image-based algorithm. Apart from the electrodes, the head mesh can be generated purely by image-based meshing, as well as it is possible to create it solely using the surface-based approach. Compared to surface-based meshing approaches where the mesh is generated from intermediate surfaces that describe the boundaries of the mesh and its individual sub-compartments, image-based approaches create the volume mesh from a labeled image and determine the boundaries of structures of different labels via a bisection algorithm. In our case, which combines image-based and surface-based meshing, the resulting mesh includes sub-compartments for every label found in the image and for every input surface.

We created a C++ tool based on the *mesh_hybrid_mesh_domain* example of the CGAL library. The tool combines the CGAL domain classes *Labeled_image_mesh_domain_3* and *Polyhedral_mesh_domain_with_features_3* into a single hybrid domain to simultaneously employ an image-based meshing together with a feature-preserving, surface-based meshing. As input, the tool requires an ANALYZE label image of the subject comprising only the structures, for which the image-based meshing should be used, as well as the OFF surface descriptions of the electrodes, the scalp and any structure, for which the surface-based meshing approach is favored. The feature edges of the electrodes are only preserved if the scalp is provided as a surface too.

To create the boundary surfaces for the surface-based volume meshing, we suggest a three-stage process. The initial boundary surface descriptions are generated from the segmentation label image by employing the Contour filter in ParaView, which is based on the Marching-cubes algorithm. Second, to take full advantage of the accurate preservation of boundaries of the surface-based meshing, the coarse output surfaces of the Contour filter must be smoothed in Meshlab using the Taubin smoothing algorithm (λ = 0.5, μ = − 0.53, #*smoothing steps* = 50). The smoothed scalp surface as a result of the electrode placement procedure does not require additional smoothing. Finally, the quality of the smoothed surface meshes must be improved by clearing defects (e.g. self-intersecting triangles) using the MeshFix tool (v.2.1) (30) and by employing a custom tool leveraging the isotropic remeshing functionality of CGAL’s *Polygon_mesh_processing* class.

To minimize the deviations from the boundaries of the labeled structures during the image-based meshing a small tolerance parameter (10 ^−6^≅0.00044 *mm* at 1 mm voxel size) for the bisection algorithm is used. Following the initial mesh generation, four optimizations can be optionally enabled. Two global optimizers (*Optimized Delaunay Triangulation smoother*, *Lloyd smoother*) minimize the total mesh energy. Two local optimizers improve the dihedral angles of the worst cells in the mesh or eliminate triangles with a poor radius-edge ratio, so-called slivers, respectively. For further information on these four optimizers, please refer to the CGAL documentation (https://doc.cgal.org/latest/Mesh_3/group__PkgMesh__3Functions.htm). We use the API of GMSH v.4.3 (31) to export the resulting volume mesh to the GMSH file format version 2.

The generated mesh is subsequently converted to the OpenFOAM format and optimized for the later computations using the OpenFOAM utilities *gmshToFoam, transformPoints,* and *renumberMesh* (details in Fig.1).

#### 2.1.4 Conductivity values

We use the OpenFOAM *setFields* tool to uniformly set a distinct isotropic tensor value for all elements of each sub-compartment of the mesh. This value is computed as the product of the unitary matrix and the corresponding scalar conductivity value:

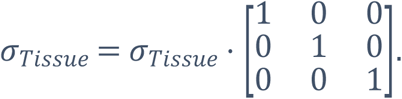

To incorporate anisotropic conductivity information of the white matter, we adopted the volume-constraint method (32). This approach assumes a shared principal direction between a diffusion tensor and its corresponding conductivity tensor but different eigenvalues representing a fixed anisotropy ratio between the principal and auxiliary directions. The calculation of the eigenvalues is based on the scalar conductivity value of the white matter *σ*_*WM*_, an anisotropy ratio of 1:10 and must satisfy the conditions 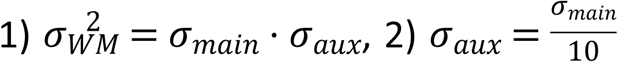 to ensure that no unreasonable conductivity values are estimated. The conductivity tensor is determined by the both-sided multiplication of the matrix *S* of the eigenvectors of the diffusion tensor and a diagonal matrix *σ*_*T*_ = *S* · *diag*(*σ*_*main*_, *σ*_*aux*_,*σ*_*aux*_) · *S*^*T*^.

The DWI data are preprocessed using MRtrix 3 (20). First, the signal to noise ratio of the DWI data is improved (*dwidenoise* (33), (34)). Subsequently, artifacts due to eddy currents and due to motion are corrected (*dwipreproc* (35), (36)). For skull-stripping, a binary mask of the intracranial tissue is generated (*dwi2mask* (37)). Tensor estimation is realized through *dwi2tensor* (38). The resulting tensor image is used to compute the fractional anisotropy (FA) map (*tensor2metric* (39), (40)). Both the FA map as well as the tensor image are cleaned from possible NaN values using *fslmaths*. The FA map is registered to the T1-weighted brain image of the subject linearly using FSL *FLIRT* (41), (42) and non-linearly with FSL *FNIRT* (43), (36). The calculated transformations are utilized to co-register the diffusion tensor image using the tool *vecreg* which preserves the relative orientation of the tensors upon transformation. The computation of the conductivity tensors is implemented as a ParaView plugin. They are subsequently transferred to the OpenFOAM mesh of the respective head model in ParaView and finally exported in the OpenFOAM field format using another custom plugin. The field values are transferred to the already prepared field of isotropic conductivity tensors, overwriting the values of the white matter compartment.

#### 2.1.5 Boundary conditions

A Dirichlet Boundary condition for the electrical potential of +/- 5 V is assigned to the outer boundaries of the anode and cathode respectively, regardless of the desired current strength. During post-processing, the electrical field strength magnitude is corrected according to the actual current density integrated at the contact surfaces of both electrodes with the scalp. The outer boundaries of the electrodes are, thus, modeled as equipotential surfaces. Since the surrounding air is not explicitly modeled and virtually acts as an insulator, a zero gradient Neumann boundary condition is applied for the electrical potential at the scalp surface.

### 2.2 Solving the electrostatic problem

The electrical field strength E and the field of the electrical current density J are computed according to the quasi-static form of Maxwell’s equations which provide a sufficient approximation for tDCS, tACS, and tRNS (44). Their solution is derived by our solver application using the OpenFOAM API.

#### 2.2.1 Quasi-static form of Maxwell’s equations

The electrical potential field *ϕ* induced by the electrodes subject to the conductivity *σ* of the volume conductor is described by Laplace’s equation ∇(*σ* ⋅ ∇ ⋅ *ϕ*) = 0. *E* is obtained by the component-wise partial derivation of *ϕ, E* =− ∇ ⋅ *ϕ*. A linear relationship between E and J by *σ* exists as *J* = *σ* · *E*.

#### 2.2.2 The solver application

Our solver application computes the electrical current density *J* and the electrical field strength *E* using the finite-volume method (FVM).

First, *ϕ* is computed using a Gauss discretization scheme with linear interpolation for the Laplace operator at a residual of 10 ^−6^. The solution is iterated to correct for non-orthogonality in the mesh until the residual of the whole solution falls below 10 ^−5^. Next, the gradient field *E* of *ϕ* is determined using the least-squares gradient scheme. *J* is the product of *E* and the electrical conductivity *σ*.

Finally, *E* and *J* are scaled by the ratio 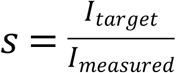 of the user-defined input current strength *I_target_* and the actual current strength *I_measured_* as determined by the summation of the current density across the surface area where the electrodes contact with the scalp surface.

### 2.3 Visualization

Post-processing is handled by ParaView for which OpenFOAM provides a plugin to read the results. All figures relating to simulation results have been created in ParaView.

## 3 Results

We demonstrate a 3-step verification attempt of the proposed workflow. First, our solver application was tested using an analytically verifiable, 3-layered sphere model (45). Second, we utilized two reference head models, which were generated in SimNIBS 3.0, to conduct tDCS simulations in both, OpenFOAM and SimNIBS to compare the results using identical head models. While other simulation pipelines are equally valid for comparing purposes, we chose the SimNIBS pipeline because of the availability of test datasets. Finally, both head models were reproduced from their original MR image, respectively, using our modeling workflow and a tDCS simulation in OpenFOAM was performed. The simulation result obtained using these custom head models were compared to the previous results.

In addition, we demonstrate the capability to model anisotropic conductivity, the modeling of alternative electrode shapes, namely small circular electrodes that are used for Laplacian-tDCS, as well as the inclusion of irregular structures, lesions of the white matter, into the head model.

### 3.1 Analytical test case: 3-layer sphere model

We implemented the analytical solution to the tES problem with point electrodes in a 3-layered sphere according to (45), (46) in Python and contrasted the result with the numerical simulation results obtained by our solver application. Table 1 provides an overview of the model parameters. Since the analytical case assumes a point electrode, which cannot be modeled in OpenFOAM, we simulated a 2 mm smaller sphere in OpenFOAM and used the analytical values greater than the 85^th^ percentile of the boundary of this sphere as the Dirichlet boundary condition of the numerical simulation. The spherical domain consisted of 15.1 M. tetrahedra.

We found an overall agreement in the distribution of the electrical potential between the analytical (Fig. 3A) and numerical solution resulting in a normalized root-mean-square deviation of only 2.1% across the entire domain. The norm of the numerically calculated electrical potential tends to decline slightly stronger as compared to the analytically derived potential (Fig. 3B).

**Fig 3.**
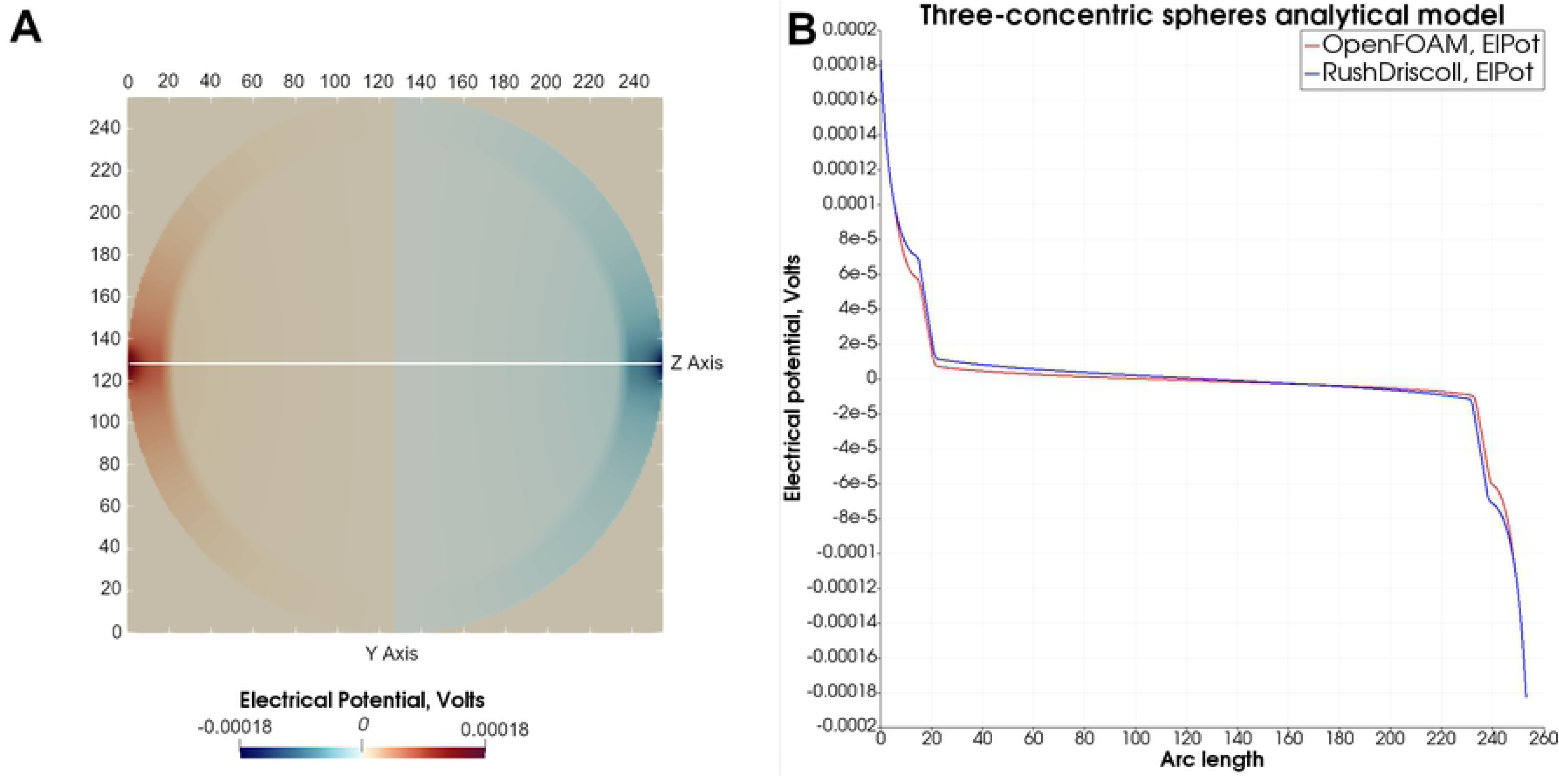
Analytical 3-layered sphere model. **A)** Center slice of the analytical result field, illustrating the distribution of the electrical potential between the two opposing point electrodes. **B)** Comparison of the electrical potential calculated analytically according to (45) (blue graph) with the numerical solution derived by OpenFOAM (red graph).

### 3.2 Comparison to SimNIBS

Our workflow was evaluated using the *Almi5* and *Ernie* test datasets from SimNIBS. Simulation results were compared to that of SimNIBS.

#### 3.2.1 Comparison of the solver application using the same head model

We utilized SimNIBS 3.0 to create the head models of the two test data sets from their T1- and T2-weighted imaging data. Each head model included the tissues skin, skull, CSF, GM, and WM. Compartments representing air were treated as a perfect insulator and were thus not part of the computational domain. For each head model, we tested three electrode setups, a bihemispheric setup over the primary motor cortices of both hemispheres, referred to as the dual setup, (10-20 positions: C3 and C4), an anodal setup (10-20 positions: C3, right supraorbital close to Fp2) and an occipital setup (10-20 positions: Cz, Oz) (Fig. 4). In all cases, square-shaped electrodes with 25 cm^2^ dimensions were modeled as a complete electrode model with equipotential surfaces at the outer boundaries. Isotropic conductivities were adopted from the SimNIBS GUI (Table 2). The input current strength was 2 mA.

**Fig 4.**
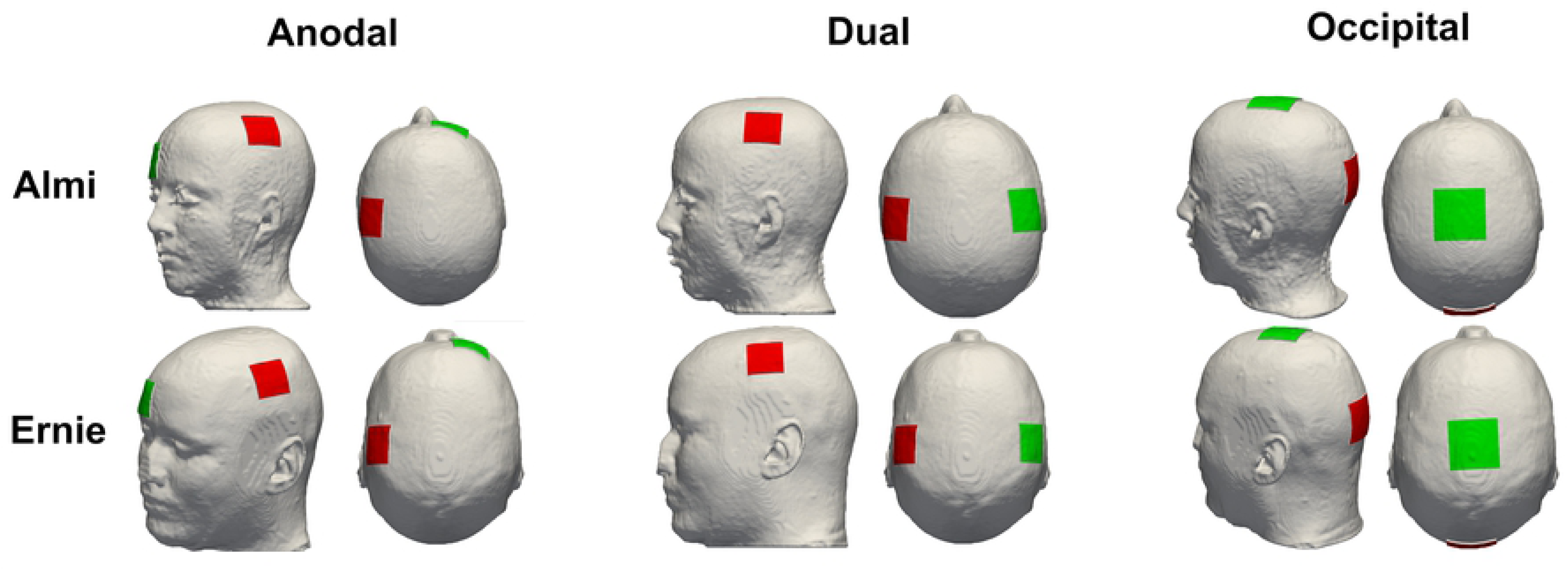
Electrode configuration. Display of the anodal, dual and occipital electrode configuration of both head models, Almi and Ernie, used for comparison with SimNIBS.

**Table 1.** Parameters of the 3-layerd spherical head model.

|  | Layer 1<br>(Scalp) | Layer 2<br>(Skull) | Layer 3<br>(Brain) |
| --- | --- | --- | --- |
| **Radii (mm)** | 92 (90 in the numerical simulation) | 85 | 80 |
| **Conductivity (S/m)** | 0.465 | 0.01 | 0.33 |

**Table 2.**
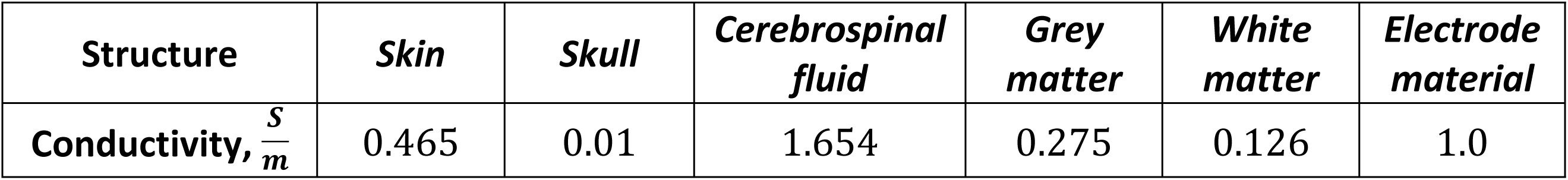
Scalar conductivity values. The conductivity values of the different head model compartments used for tDCS simulations both in OpenFOAM and SimNIBS.

| Structure | *Skin* | *Skull* | *Cerebrospinal fluid* | *Grey matter* | *White matter* | *Electrode material* |
| --- | --- | --- | --- | --- | --- | --- |
| Conductivity, $\frac{S}{m}$ | 0.465 | 0.01 | 1.654 | 0.275 | 0.126 | 1.0 |

Visual comparison of the computed electrical field strength to the field obtained by SimNIBS revealed a comparable field pattern with hotspots at the same locations across both head models and all electrode montages (Fig. 5). The magnitude of the electrical field within the gray matter mesh compartment was on average higher in our results across both models and all electrode montages (Table 3). See tables 4 & 5 and Figs. 6 – 9 for a more detailed overview of the relative difference in magnitude of the electrical field strength as well as the angle difference across all conditions. The deviation in direction was more pronounced in the area of the gray matter mesh compartment underneath the electrodes in all cases with a 99^th^ percentile peak value in angle difference of 40.56° in the occipital electrode configuration of the Almi test case. We contrasted the magnitude of the electrical field along a sampling line between the respective electrode pair of each condition through the entire head model (Figs. 10 & 11). This assessment confirmed that our simulation slightly overestimates the magnitude of the electrical field in the intracranial compartments. Interestingly, this trend reverses for skin and skull, where a small underestimation can be observed. No major difference between head models and electrode conditions was noticeable. The simulation time was approximately 4 minutes in all cases.

**Fig 5.**
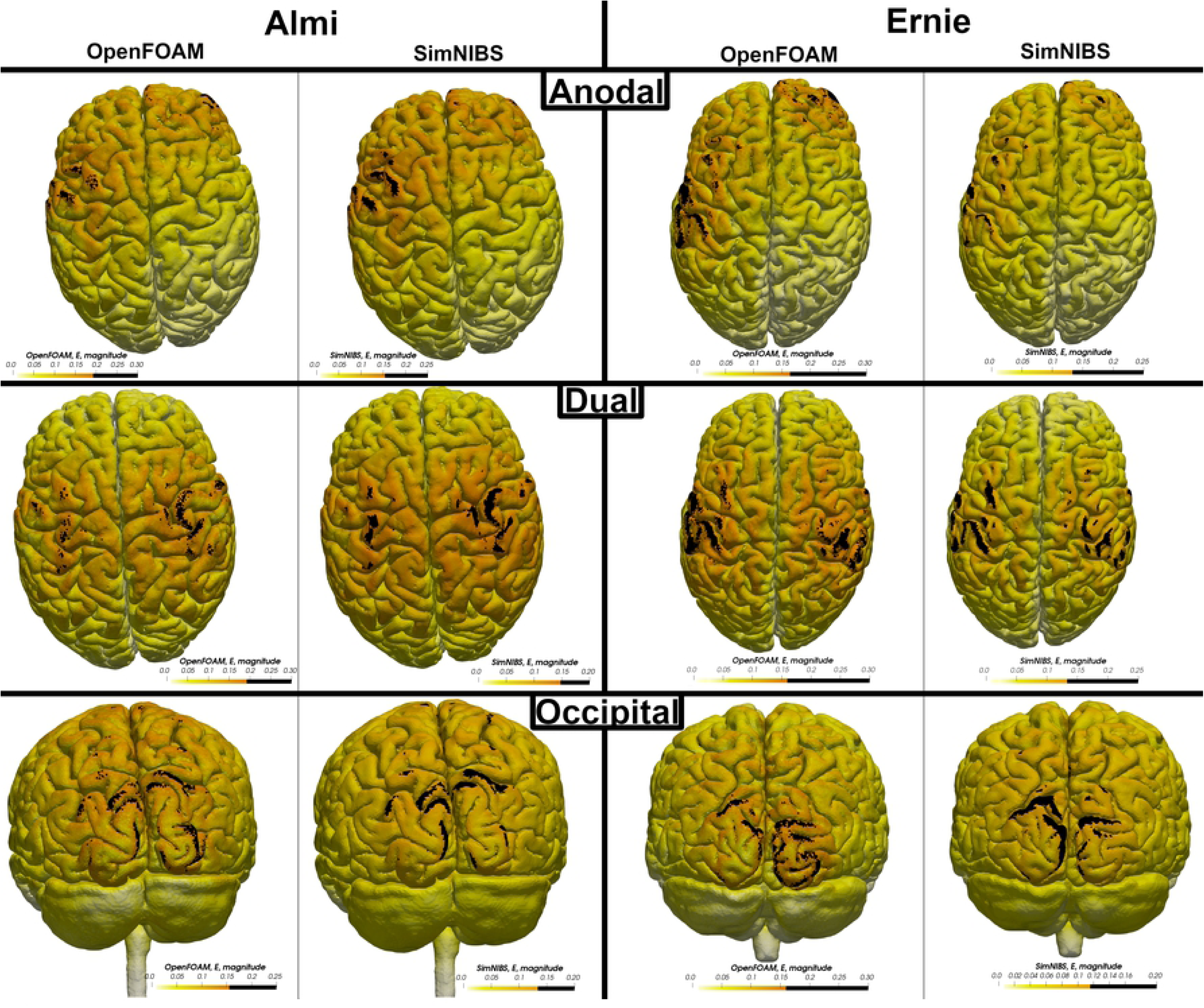
Electrical field pattern. Comparison of distribution pattern of the electrical field strength in both head models and all electrode montages between the OpenFOAM result and the SimNIBS result. Areas above the 90^th^ percentile of the electrical field strength are defined as hotspots and marked in black.

**Fig 6.**
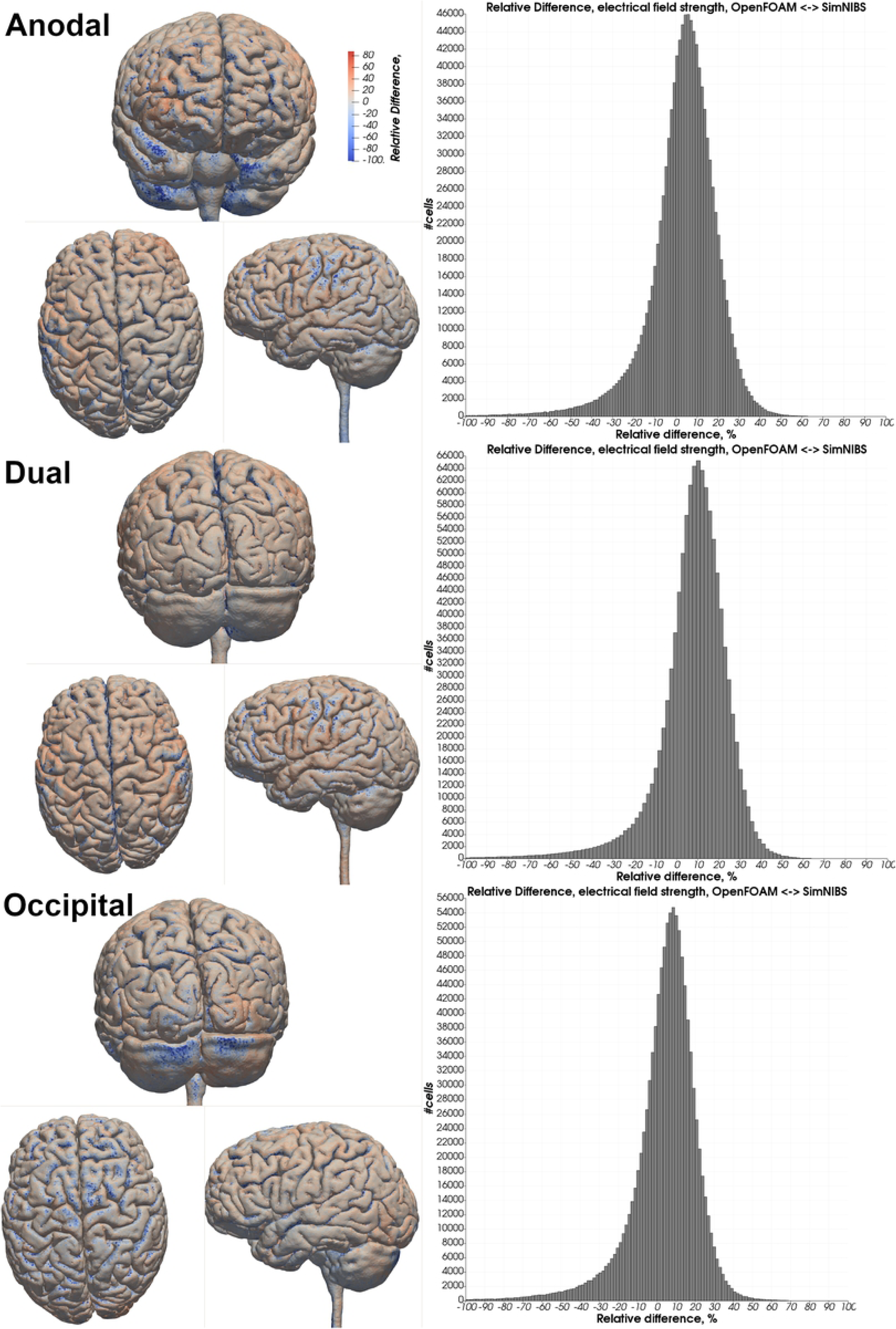
Relative difference in the electrical field strength magnitude - Almi. Heatmap of the relative difference in the magnitude of the electrical field strength between OpenFOAM and SimNIBS in all three electrode configurations. A red color indicates a higher electrical field strength in the OpenFOAM result whereas blue indicates a higher value in the SimNIBS result. Histograms depict differences in percent of all tetrahedra within the gray matter mesh compartment.

**Fig 7.**
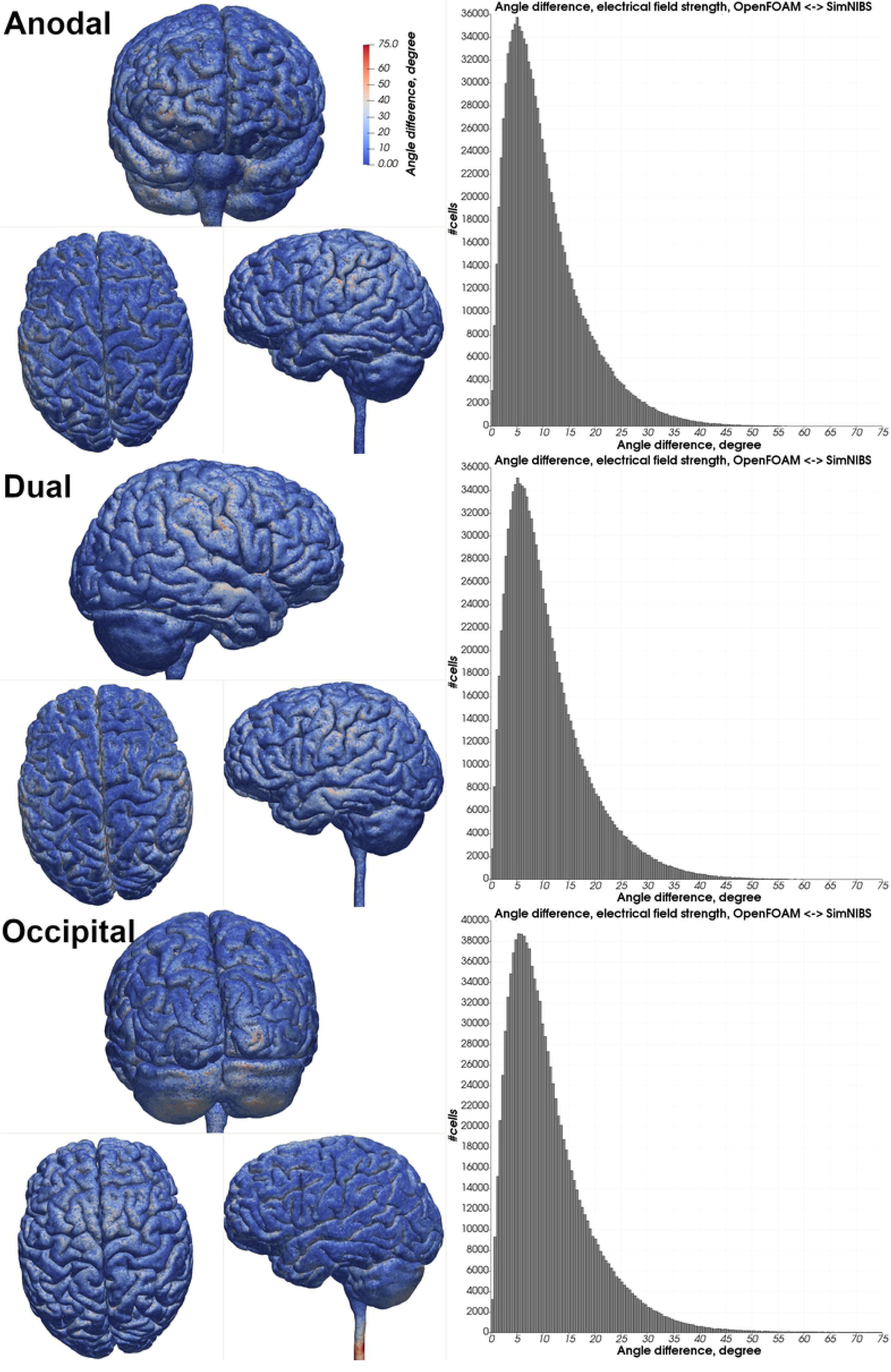
Angle difference in the electrical field strength - Almi. Heatmap of the angle difference of the electrical field strength between OpenFOAM and SimNIBS of all electrode configurations. Histograms depict angle differences in degrees of all tetrahedra within the gray matter mesh compartment.

**Fig 8.**
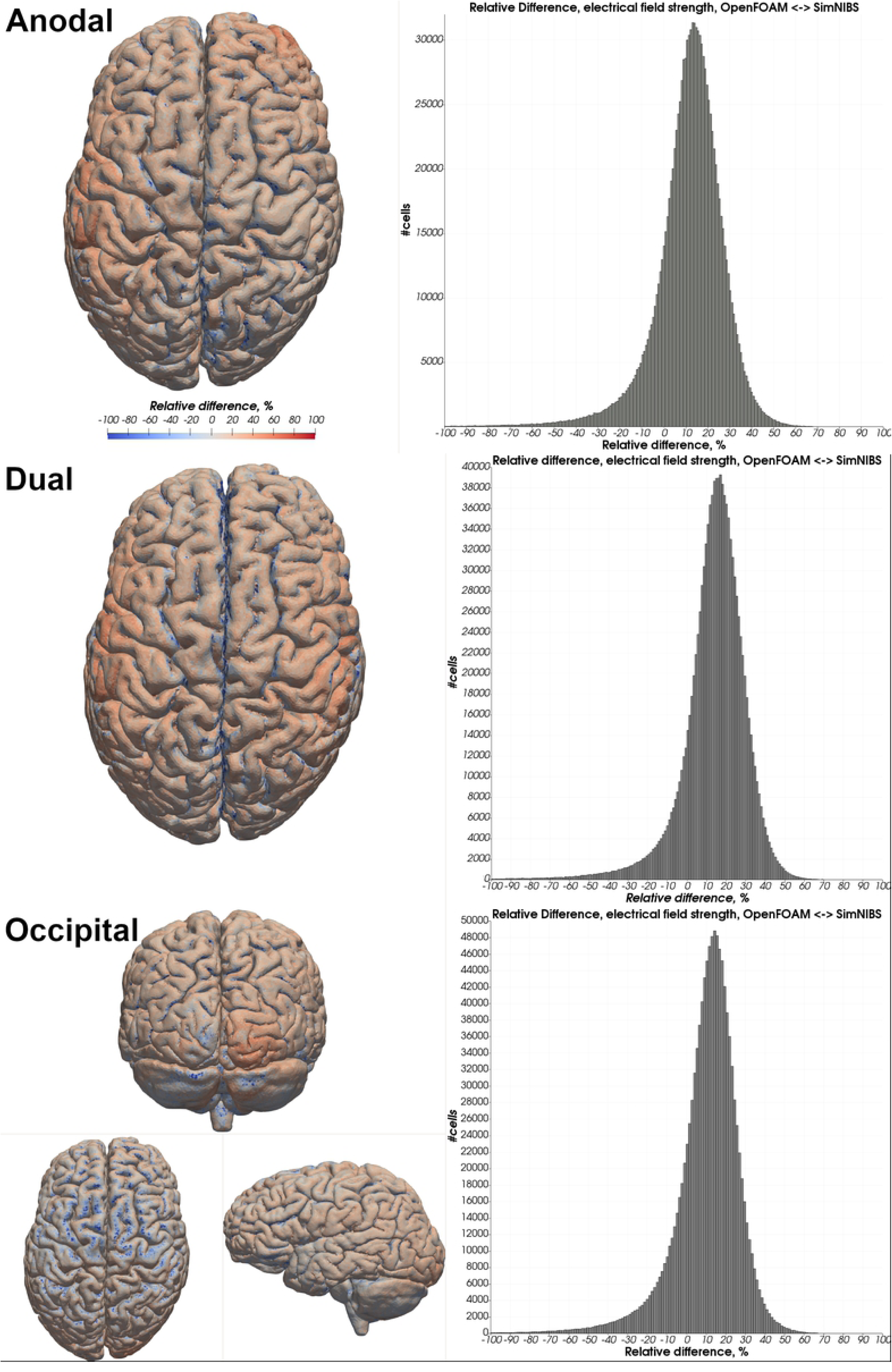
Relative difference in electrical field strength magnitude - Ernie. Heatmap of the relative difference in the magnitude of the electrical field strength between OpenFOAM and SimNIBS in all three electrode configurations. A red color indicates a higher electrical field strength in the

**Fig 9.**
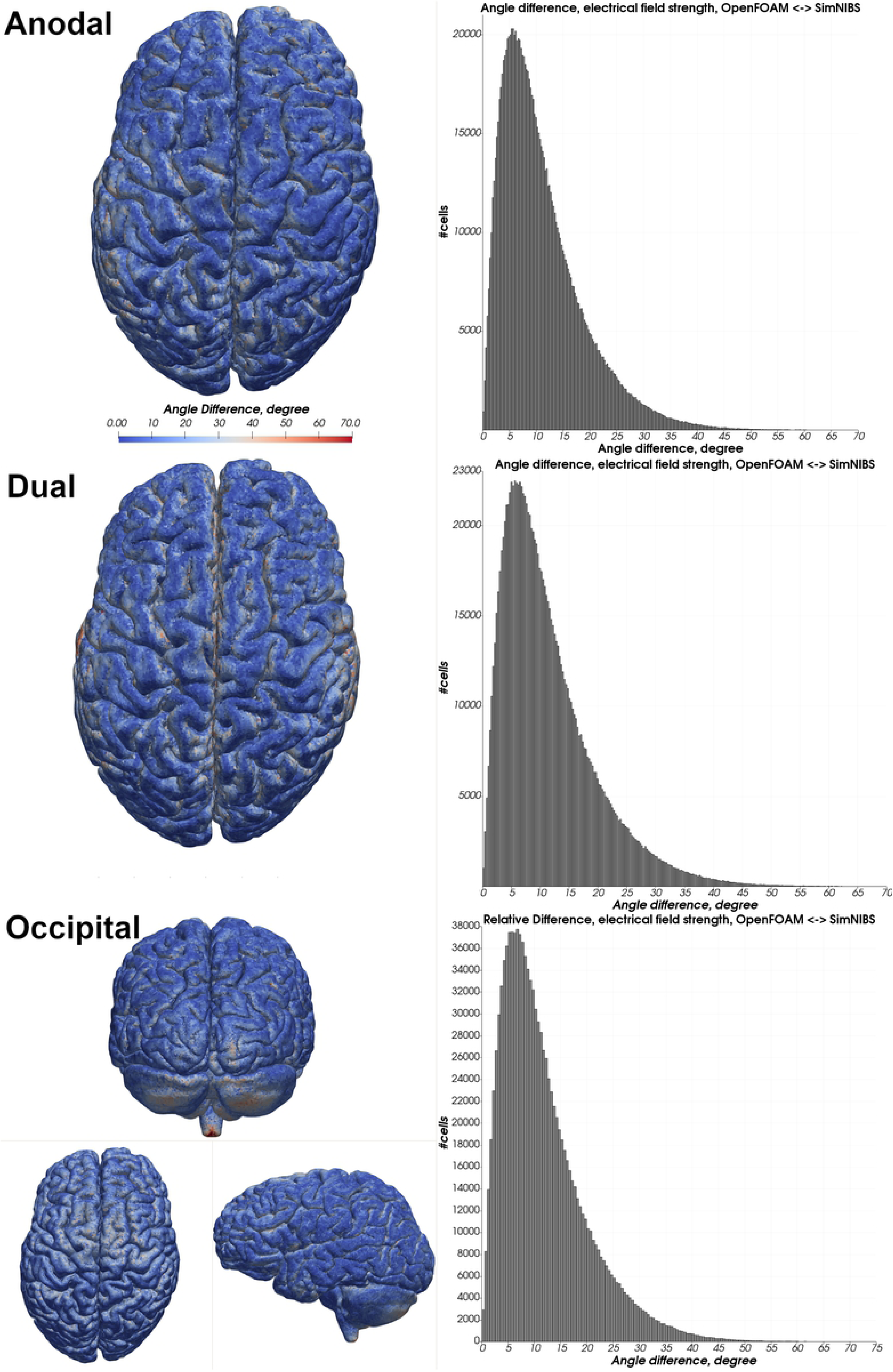
Angle difference in electrical field strength - Ernie. Heatmap of the angle difference of the electrical field strength between OpenFOAM and SimNIBS of all electrode configurations. Histograms depict angle differences in degrees of all tetrahedra within the gray matter mesh compartment.

**Fig 10.**
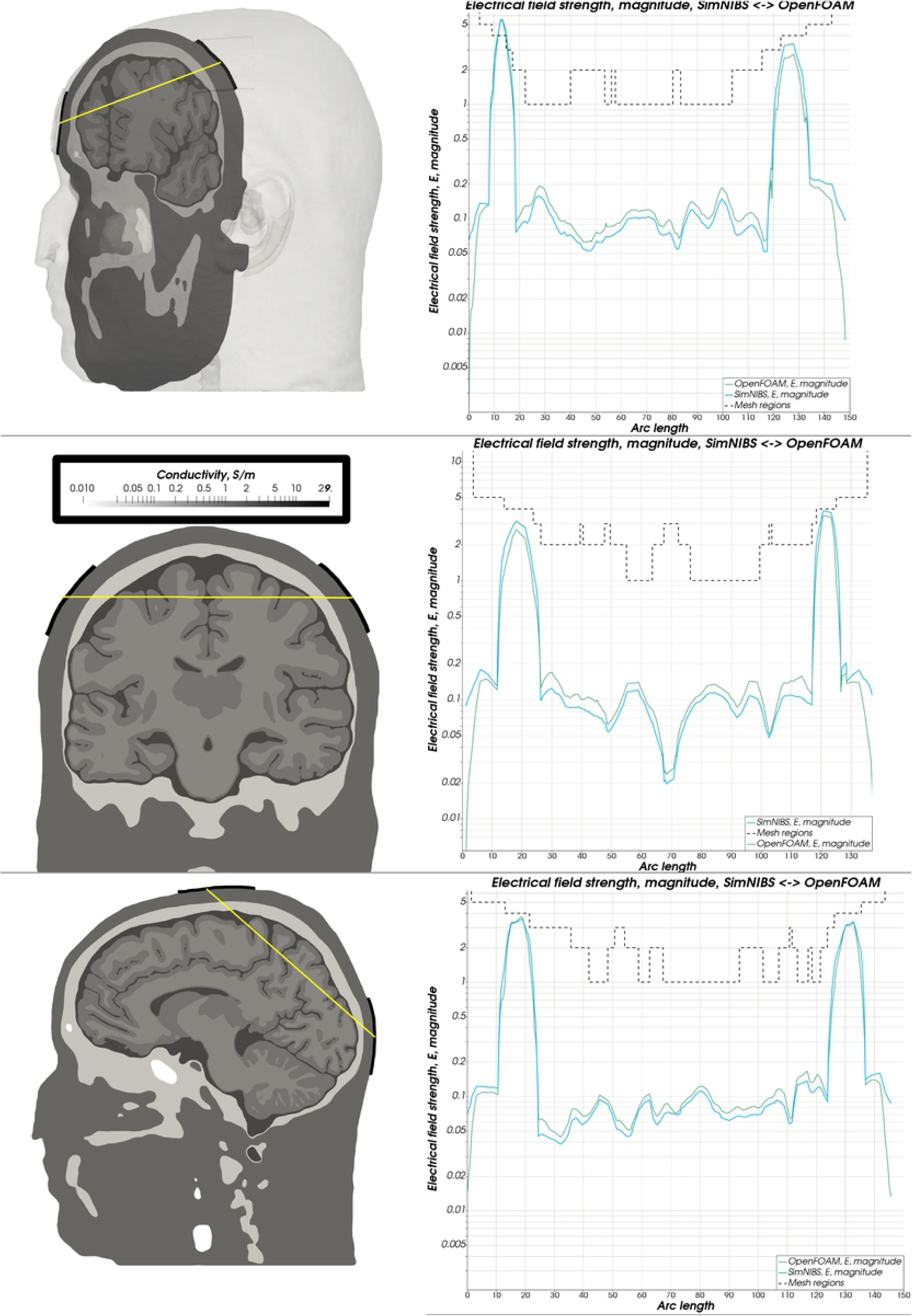
Electrical field magnitude - Almi. Comparison of the magnitude of the electrical field strength along a sampling line between both electrodes between OpenFOAM (green) and SimNIBS (blue). A dashed line depicts the mesh regions with distinct conductivity values.

**Fig 11.**
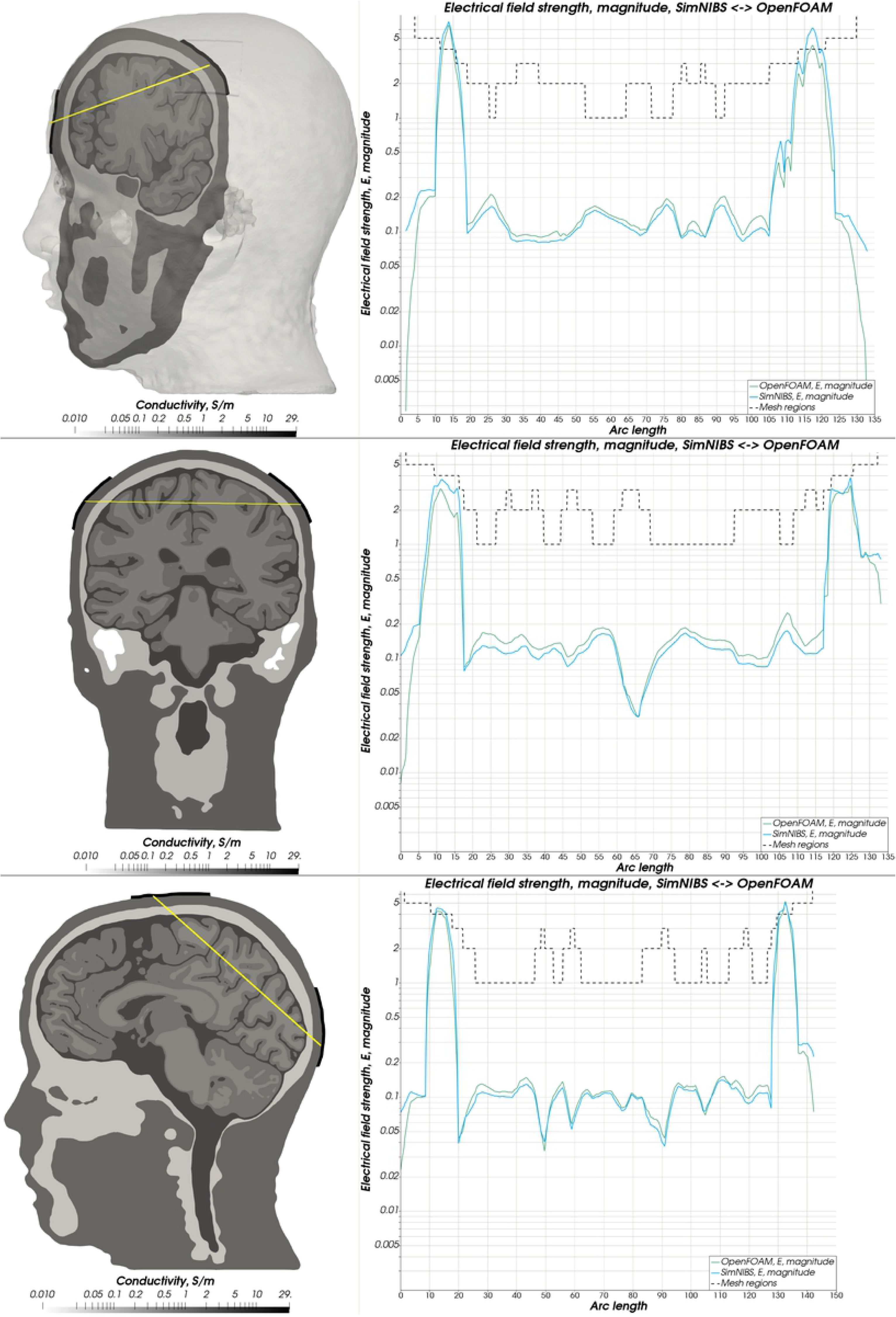
Electrical field magnitude - Ernie. Comparison of the magnitude of the electrical field strength along a sampling line between both electrodes between OpenFOAM (green) and SimNIBS (blue). A dashed line depicts the mesh regions with distinct conductivity values.

**Table 3.**
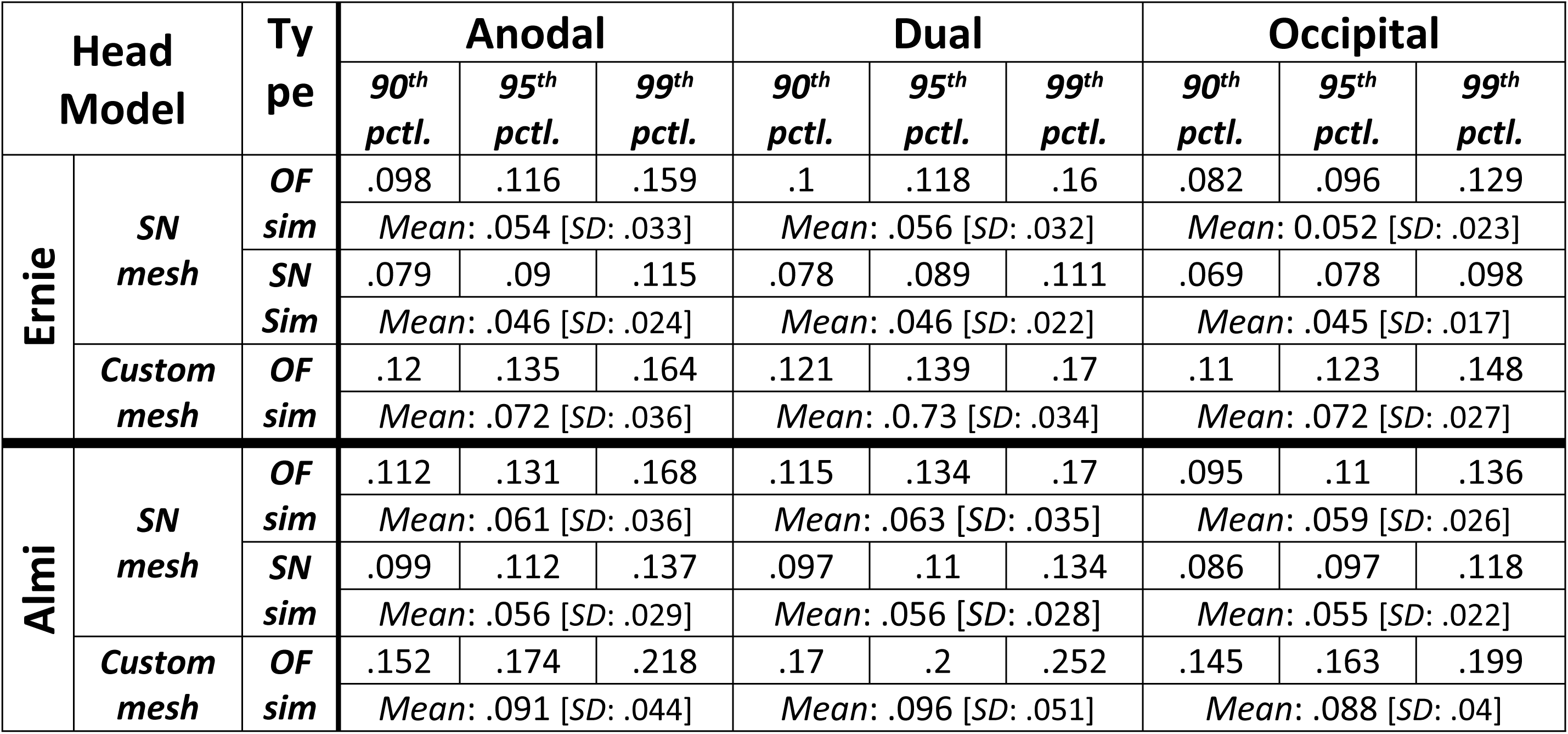
Comparison electrical field strength (SimNIBS [SN] vs. OpenFOAM [OF]). Comparison of the 90th, 95th, 99th percentile as well as the average magnitude of the electrical field strength in V/m within the gray matter mesh compartment of the head model generated by SimNIBS (“SN mesh”) and the head model generated using our head modeling pipeline (“custom mesh”). For the SN mesh, simulations in both SimNIBS and OpenFOAM have been performed. The custom mesh was only used in our simulation environment, not in SimNIBS.

**Table 4.** Absolute value of the relative difference of the electrical field strength (SimNIBS vs OpenFOAM). Comparison (in percent, 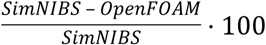) of the mean and peak percentile 100) of the mean and peak percentile absolute value of the relative difference of the simulation results computed by SimNIBS and OpenFOAM within the gray matter compartment of the reference meshes.

|  | Anodal |  |  | Dual |  |  | Occipital |  |  |
| --- | --- | --- | --- | --- | --- | --- | --- | --- | --- |
|  | 90<sup>th</sup><br>pctl. | 95<sup>th</sup><br>pctl. | 99<sup>th</sup><br>pctl. | 90<sup>th</sup><br>pctl. | 95<sup>th</sup><br>pctl. | 99<sup>th</sup><br>pctl. | 90<sup>th</sup><br>pctl. | 95<sup>th</sup><br>pctl. | 99<sup>th</sup><br>pctl. |
| **Ernie** | 29.85 % | 35.8 % | 55.4 % | 32.81 % | 38.28 % | 55.8 % | 30.32 % | 35.8 % | 51.47 % |
|  | Mean: 16.23 % [SD: 12.95 %] |  |  | Mean: 18.44 % [SD: 14.87 %] |  |  | Mean: 16.31 % [SD: 12.32 %] |  |  |
| **Almi** | 26.91 % | 34.89 % | 66.21 % | 28.2 % | 34.49 % | 63.29 % | 20.63 % | 25.34 % | 35.22 % |
|  | Mean: 13.71 % [SD: 14.37 %] |  |  | Mean: 14.9 % [SD: 15.31 %] |  |  | Mean: 10.39 % [SD: 7.52 %] |  |  |

**Table 5.** Angle difference of the electrical field strength (SimNIBS vs. OpenFOAM). Comparison of the simulation results (in degrees, 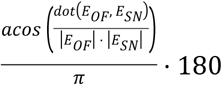) computed by SimNIBS and OpenFOAM 180) computed by SimNIBS and OpenFOAM within the gray matter compartment of the reference meshes.

|  | Anodal |  |  | Dual |  |  | Occipital |  |  |
| --- | --- | --- | --- | --- | --- | --- | --- | --- | --- |
|  | 90<sup>th</sup><br>pctl. | 95<sup>th</sup><br>pctl. | 99<sup>th</sup><br>pctl. | 90<sup>th</sup><br>pctl. | 95<sup>th</sup><br>pctl. | 99<sup>th</sup><br>pctl. | 90<sup>th</sup><br>pctl. | 95<sup>th</sup><br>pctl. | 99<sup>th</sup><br>pctl. |
| **Ernie** | 21.55° | 26.25° | 36.34° | 22.37° | 27.36° | 37.9° | 22.81° | 27.73° | 37.76° |
|  | Mean: 11.05° [SD: 7.75°] |  |  | Mean: 11.44° [SD: 8.06°] |  |  | Mean: 11.72° [SD: 8.1°] |  |  |
| **Almi** | 20.63° | 25.35° | 35.22° | 21.4° | 26.4° | 36.83° | 22.15° | 27.45° | 40.56° |
|  | Mean: 10.39° [SD: 7.52°] |  |  | Mean: 10.81° [SD: 7.81°] |  |  | Mean: 11.31° [SD: 8.65°] |  |  |

#### 3.2.2 Full workflow verification

As a next step, we reproduced the Almi5 and Ernie head models from their original T1-weighted MR data using our head modeling workflow. Their original MR data are available from SimNIBS.

Figure 12 displays the segmentation results achieved by our approach using only the T1-weighted image in comparison to SimNIBS 3.0 using the CAT12 segmentation routines on both the T1-weighted as well as the T2-weighted image of the exemplary data set “Ernie”. The computed head models were caudally more truncated. The Mesh quality (Table 6) was well suitable for OpenFOAM. The conductivity values and the three electrode montages remained unchanged. Computation times for each head model on an Intel Core i7 6700 workstation were approximately 6 hours (segmentation), 3 hours (meshing), 100 seconds (simulation).

**Fig 12.**
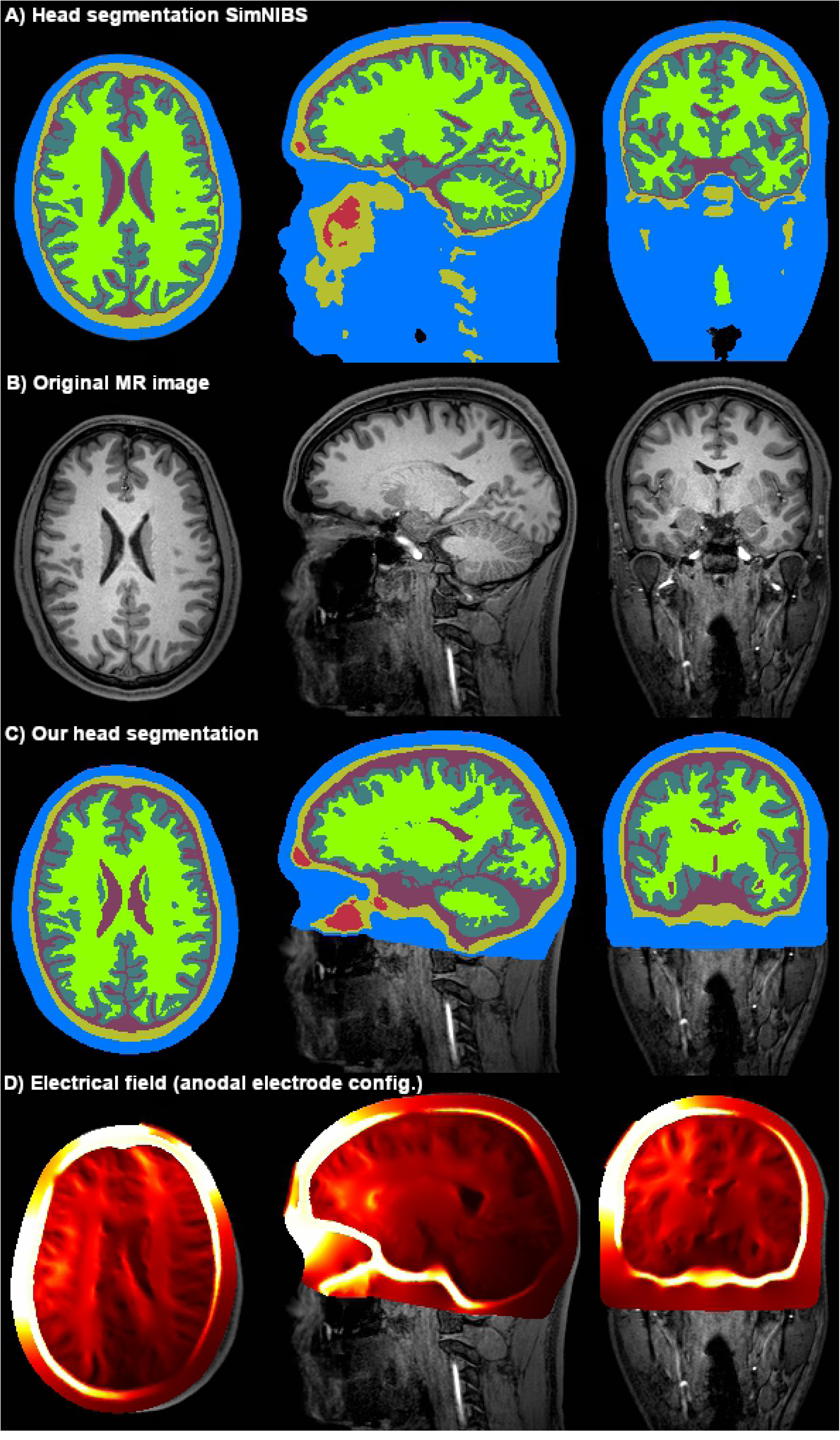
Head segmentation. Comparison of our segmentation result **(C)** of the T1-weighted MR image of the Ernie test dataset **(B)** with the SimNIBS segmentation **(A)** computed from the T1- and T2-weighted imaging. The resulting electrical field of an anodal electrode configuration using a head model generated from our segmentation is displayed in Panel **D**.

**Table 6.** Mesh characteristics. Number of cells and mesh quality metrics of our version of the Almi and Ernie head models as well as the version generated by SimNIBS. For the subsequent finite-volume-method calculation decisive characteristics are the number of mesh elements (#cells), the number of non-orthogonal faces, i.e. faces whose non-orthogonality is greater than 70°, the maximum non-orthogonality and the maximum skewness of the mesh elements.

|  | Almi (dual) |  | Ernie (dual) |  |
| --- | --- | --- | --- | --- |
|  | SimNIBS version | Our version | SimNIBS version | Our version |
| **#cells (in million)** | 4.1 | 5.2 | 4.8 | 5.5 |
| **#non-orthogonal faces** | 1090 | 239 | 2301 | 26 |
| **Max. non-orthogonality** | 93.43° | 81.04° | 89.72° | 82.12° |
| **Max. cell skewness** | 10.9 | 2.39 | 3.04 | 1.78 |

The magnitude of the resulting electrical field strength was again compared to the previous results by sampling along a sampling line between the respective electrodes through the head models. Across all conditions, the mean and percentile-peak values of the electrical field strength in the gray matter mesh compartment were slightly overestimated (“custom mesh” vs “SN mesh” in Table 3) while the field distribution (Fig. 12D) remained comparable (Fig. 13 & 14).

**Fig 13.**
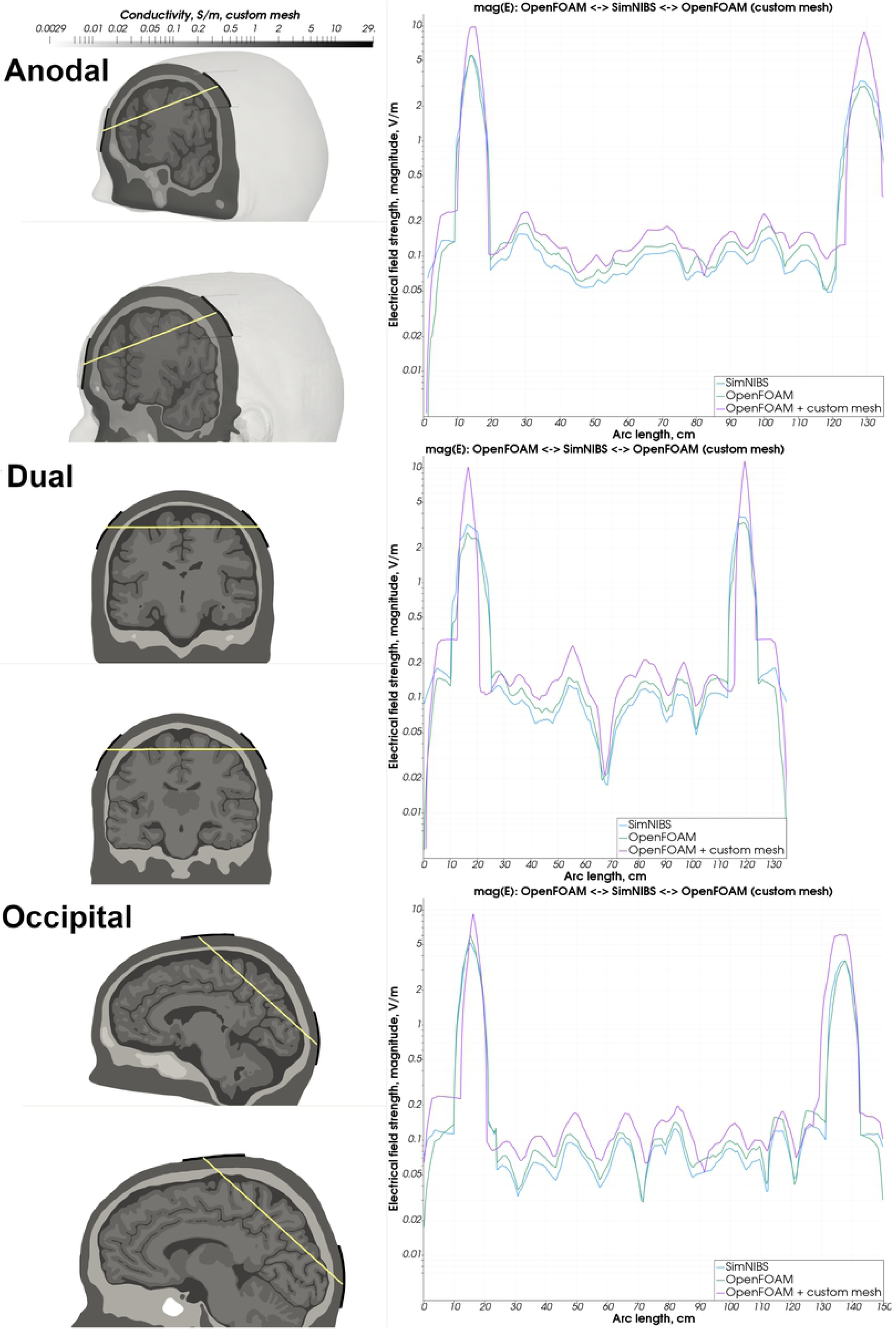
Simulation results custom mesh vs SimNIBS mesh - Ernie. Comparison of the magnitude of the electrical field strength of the simulation conducted using our custom head model (purple) and the SimNIBS generated head model (OpenFOAM result: green, SimNIBS result: blue).

**Fig 14.**
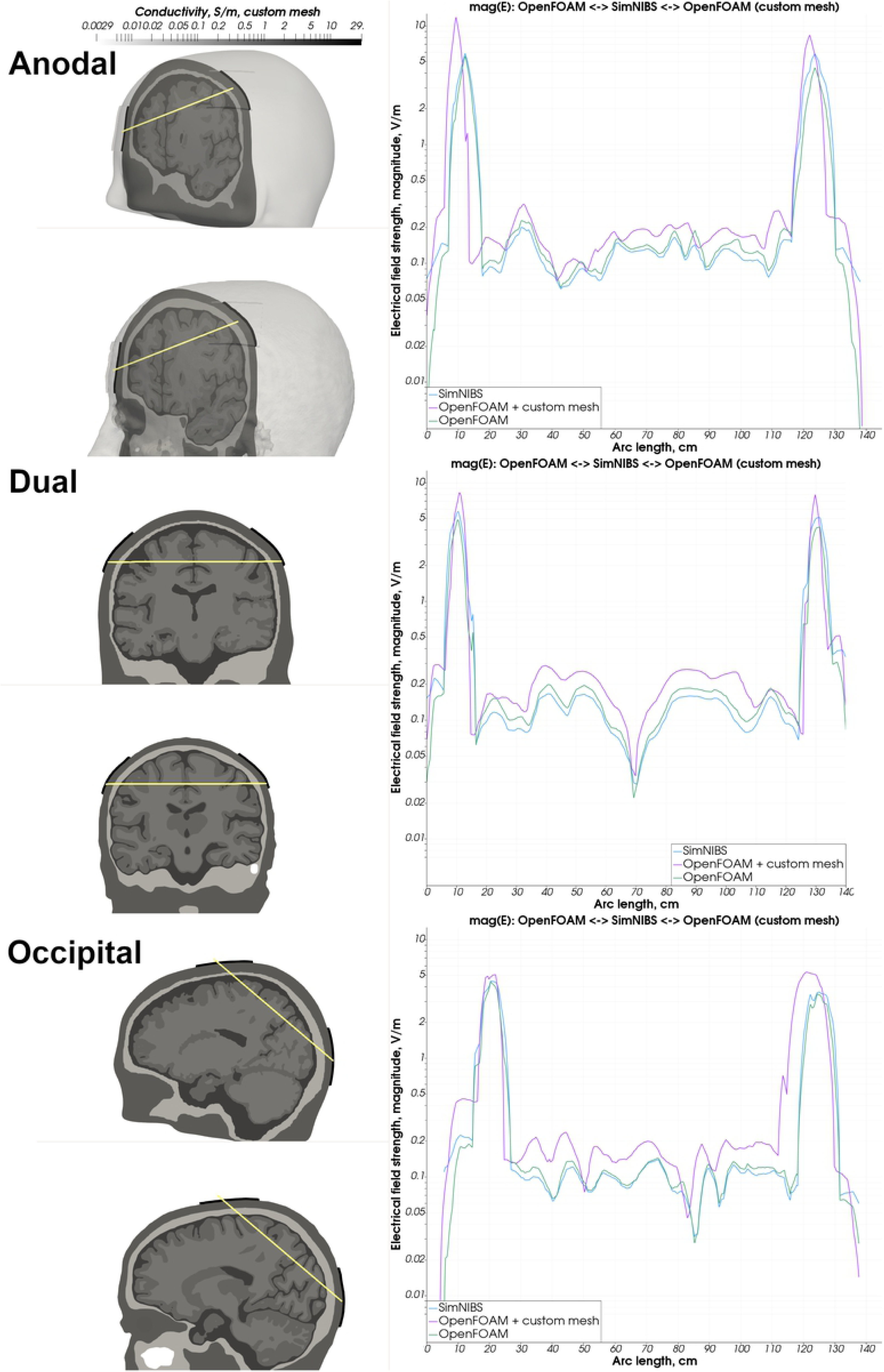
Simulation results custom mesh vs SimNIBS mesh - Almi. Comparison of the magnitude of the electrical field strength of the simulation conducted using our custom head model (purple) and the SimNIBS generated head model (OpenFOAM result: green, SimNIBS result: blue).

### 3.3 Extended capabilities

In this section, we demonstrate extended processing capabilities that can be combined with our standard workflow. We included anisotropic conductivity of the white matter in the custom Almi5 test case. Furthermore, we conducted a simulation using an alternative electrode model in the form of small circle-like electrodes in a multi-electrode setup. Finally, we demonstrate the inclusion of lesioned tissue in a head model.

#### 3.3.1 Modeling anisotropic conductivity

To model the anisotropic conductivity of white matter, the conductivity tensors from the diffusion-weighted imaging data of the Almi5 dataset were computed. In this process, we assumed a fixed ratio of 1:10 between the main and the auxiliary directions of the tensor and a conductivity of 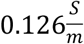 for the white matter. We assigned the same isotropic conductivity values to the individual mesh compartments as before except the white matter compartment to which we assigned the computed conductivity tensors. Refer to Fig. 15 for a depiction of the conductivity profile of the data set. We simulated the anodal electrode setup with two 5 cm x 5 cm patch-like electrodes placed over C3 and supraorbital, close to Fp2. The input current strength was set to 2 mA. Additionally, to demonstrate the image-based meshing capabilities of our meshing tool we generated the head model only using image-based meshing (except for the electrodes and the scalp to ensure the feature-preservation of the electrodes). The characteristics of the resulting mesh were as follows: 5.2 million tetrahedra, 239 non-orthogonal faces, maximum non-orthogonality of 81°, maximum skewness of 2.4.

**Fig 15.**
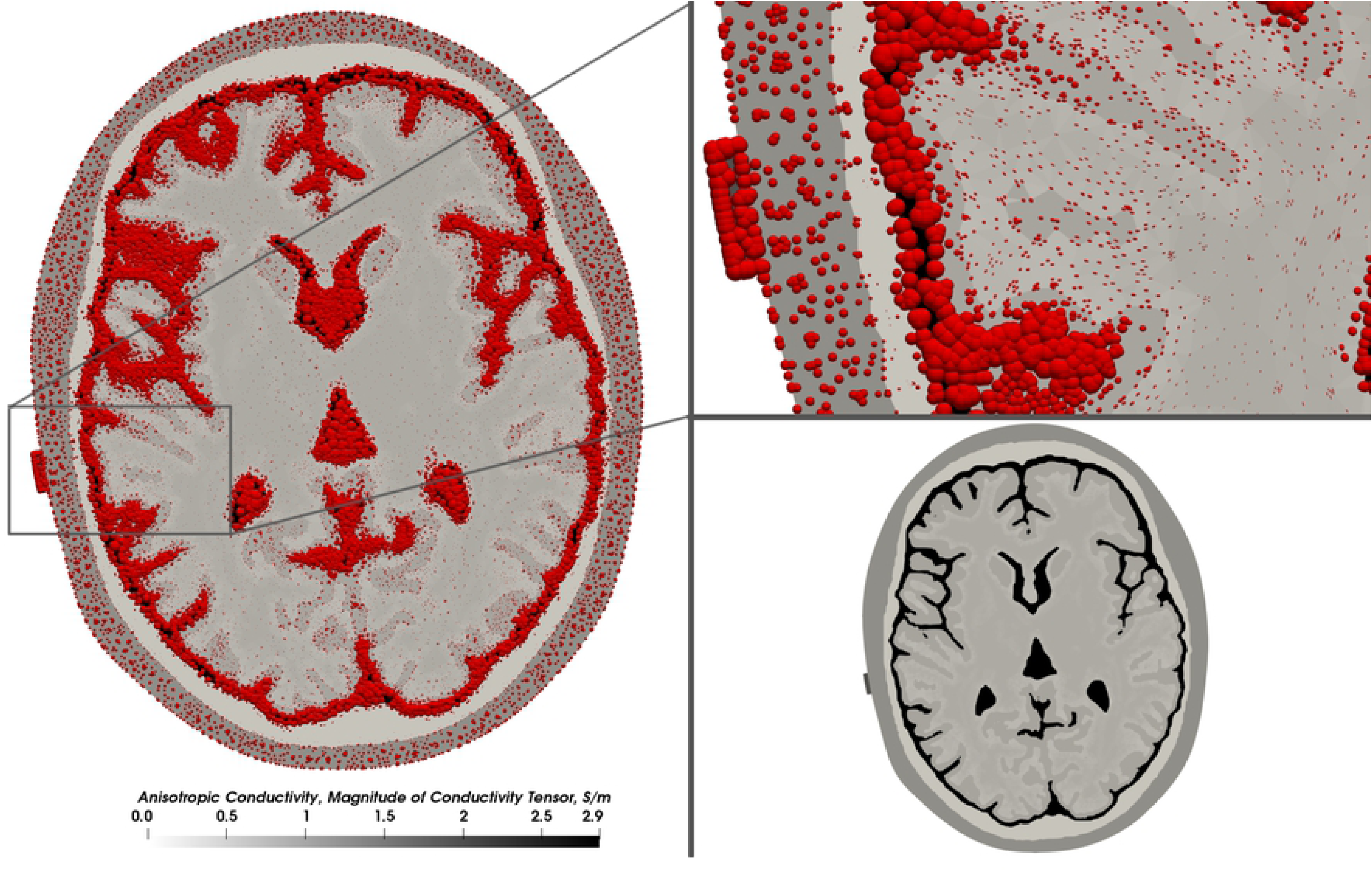
Conductivity tensors. Conductivity profile of the augmented Almi5 test case. Darker color depicts higher conductivity. Conductivity tensors are visualized using spherical tensor glyphs. Their size depicts the magnitude of the conductivity. The shape reflects the degree of anisotropy, from isotropic (ball shape) to highly anisotropic (ellipsoidal, rod-like).

We sampled the magnitude of the electrical field strength along a sampling line between both electrodes through the head model and compared the magnitude of the anisotropic test case to a version of the test case using scalar conductivity values only. The difference in the magnitude was most noticeable in the intracranial compartments, where the changes in the magnitude (both in the negative and positive direction) along the sampling line were generally higher in the anisotropic case as compared to the isotropic case (Fig. 16A). Furthermore, the area underneath the electrodes experienced higher differences both in the local field angle and field magnitude of the electrical field (Fig. 16B & 16C). The mean angle difference between the isotropic and anisotropic case within the gray matter mesh compartment was 9.4° (99^th^ percentile: 33.1°) and the mean value of the relative difference of the absolute field magnitude was 12.25 % (99^th^ percentile: 29.16%).

**Fig 16.**
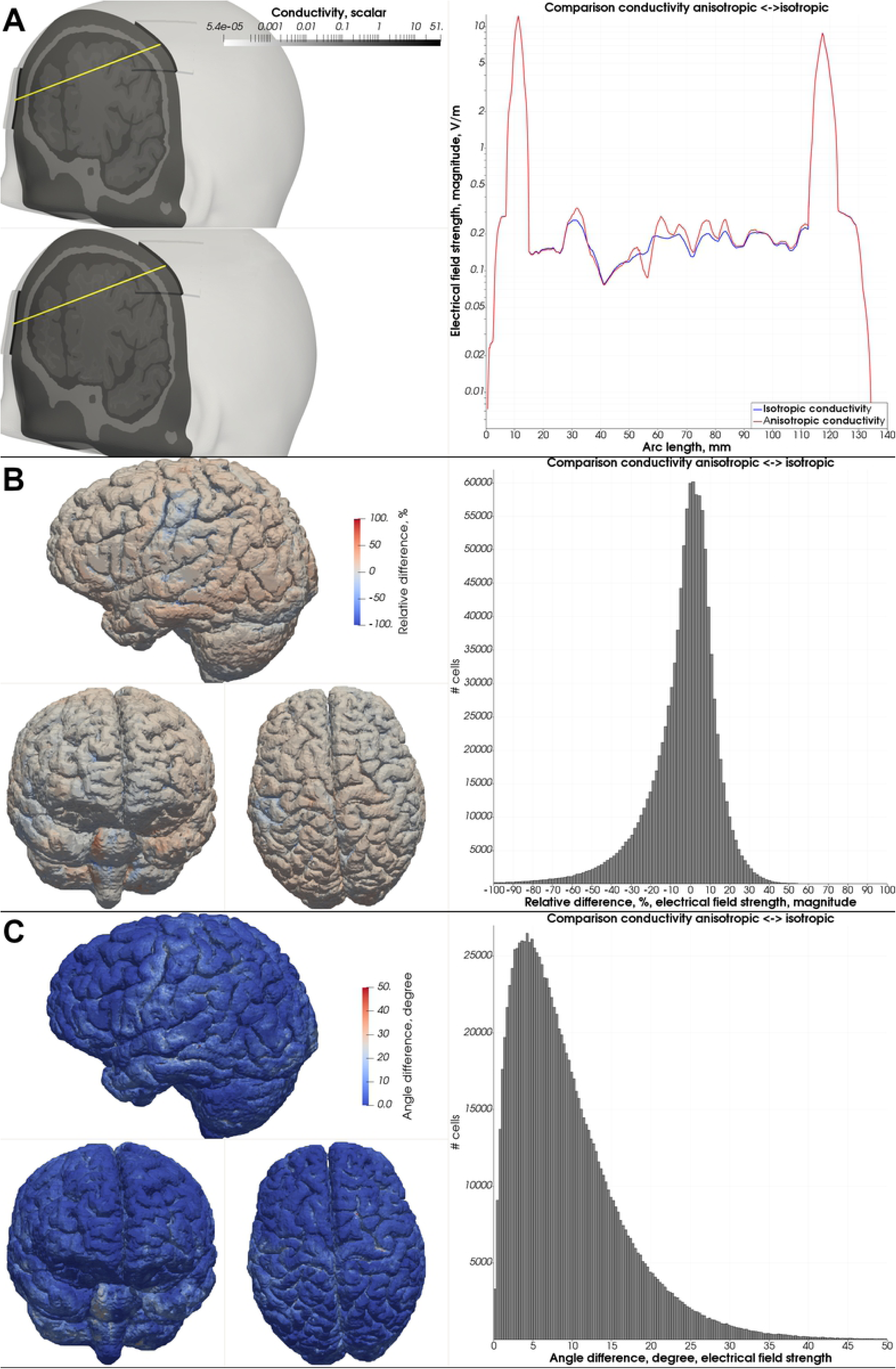
Anisotropic test case – comparison to isotropic test case. **A)** Comparison of the magnitude of the electrical field strength along the sampling line (yellow) between the custom version of the Alim5 head model with isotropic and anisotropic white matter conductivity. **B)** Relative difference

#### 3.3.2 Simulating multi-electrode tDCS

In this test case, we changed the electrode setup to a 4 x 1 multi-electrode tDCS setup with five circular electrodes with a diameter of 5 mm. The anode was positioned approximately at C3. The four cathodes were positioned in 10 cm distance from the anode in a square arrangement around the cathode. A Dirichlet boundary condition of −5 V at the four cathodes and +5 V at the central anode was defined. We set the input current strength to 2 mA. Again, the image-based meshing algorithm was used for the head model generation (surface-based only for the scalp and the electrodes). The same isotropic conductivity values as before were assigned.

The computation of the electrical field finished after 148 seconds. The resulting electrical field pattern is much more focal (Figure 17) with only a negligible fraction of the inbound current reaching the contralateral hemisphere as compared to the field induced by two large conventionally shaped electrodes as simulated before. This is an expected observation for multi-electrode tDCS montages. The average electric field strength across the cortex was reduced to 0.02 V/m. The 99^th^ percentile peak electric field strength was lowered to 0.161 V/m. A larger portion of the cortex that received non-negligible field strength is covered by a field strength of the 99^th^ percentile.

**Fig 17.**
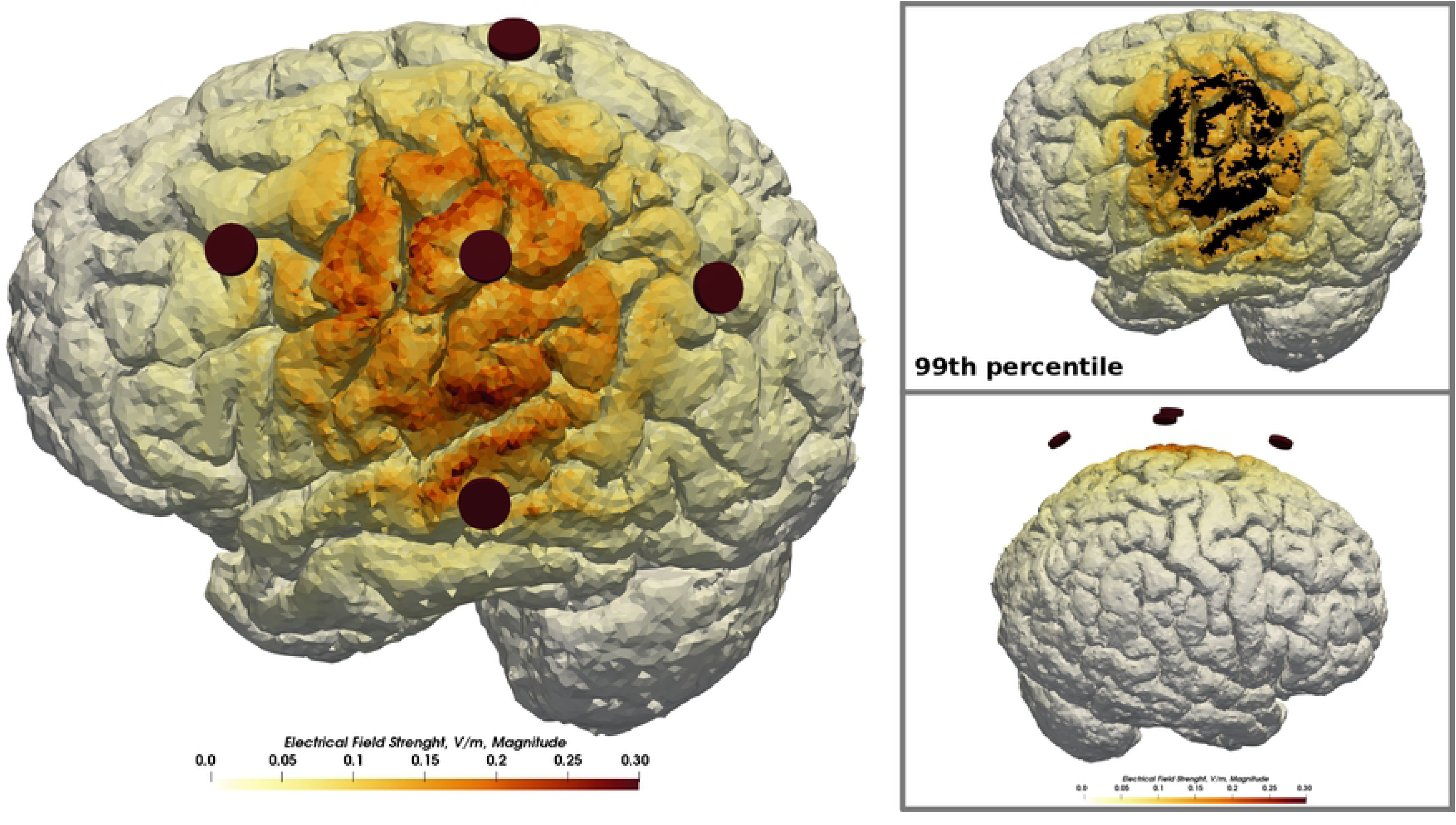
Multi-electrode test case. Exemplary extension of the standard workflow by multi-electrode tDCS. Five round electrodes with a diameter of 5 mm were positioned approximately at C3 and resulted in a much more focal field distribution than achieved with conventional square-shaped, patch electrodes.

#### 3.3.3 Inclusion of lesioned tissue

In this test case, we created a head model from the T1-weighted magnetization prepared rapid gradient echo (MRAGE) and T2-weighted fluid-attenuated inversion recovery (FLAIR) imaging data of a single subject from the local, large-scale, cross-sectional study of the Leipzig Research Centre for Civilization Diseases (LIFE) (47). Imaging parameters used for the MPRAGE image were: flip angle 9°, repetition time 2300 ms, inversion time 900 ms, echo time 2.98 ms, 1 mm isotropic resolution, acquisition time 5.1 min. The parameters of the FLAIR image were: repetition time 5000 ms, inversion time 1800 ms, echo time 395 ms, 1 mm isotropic resolution, acquisition time 7.02 min. The images were acquired on a MAGNETOM Verio scanner (Siemens, Erlangen, Germany) with a 32-channel head receive coil and a body transmit coil. The head model was generated by our robust standard segmentation workflow using the T1-weighted imaging data. Additionally, we included white-matter lesions into the head model that were segmented before using the T2-FLAIR data. Details of the white matter lesion segmentation procedure, which relied on an adapted version of the lesion-TOADS algorithm (48), can be found in (49). We employed image-based meshing for the lesioned tissue, the ventricles and the air cavities of the skull, and applied the surface-based meshing to all other structures (scalp, skull, CSF, GM, WM, electrodes).

To illustrate the robustness of our segmentation and meshing approach, we compared the generated compartments of the head mesh between our approach, SimNIBS 3.0 and ROAST 2.7.1 (Fig. 18). Our approach strongly smooths the scalp structure but maintains typical characteristics of the shape of the scalp (Fig.18A). The skull is most robustly estimated by our approach, which, however, tends to overestimate the thickness of the skull occipitally, along the superior sagittal sinus (Fig.19B), and caudally. All three approaches yield a comparable gray matter compartment (Fig.18D). SimNIBS creates the visually most complete white matter compartment (Fig. 18E). Note that we included the white matter lesions as a separate compartment only in our head model (highlighted in orange) (Fig. 18E, Fig. 19A).

**Fig 18.**
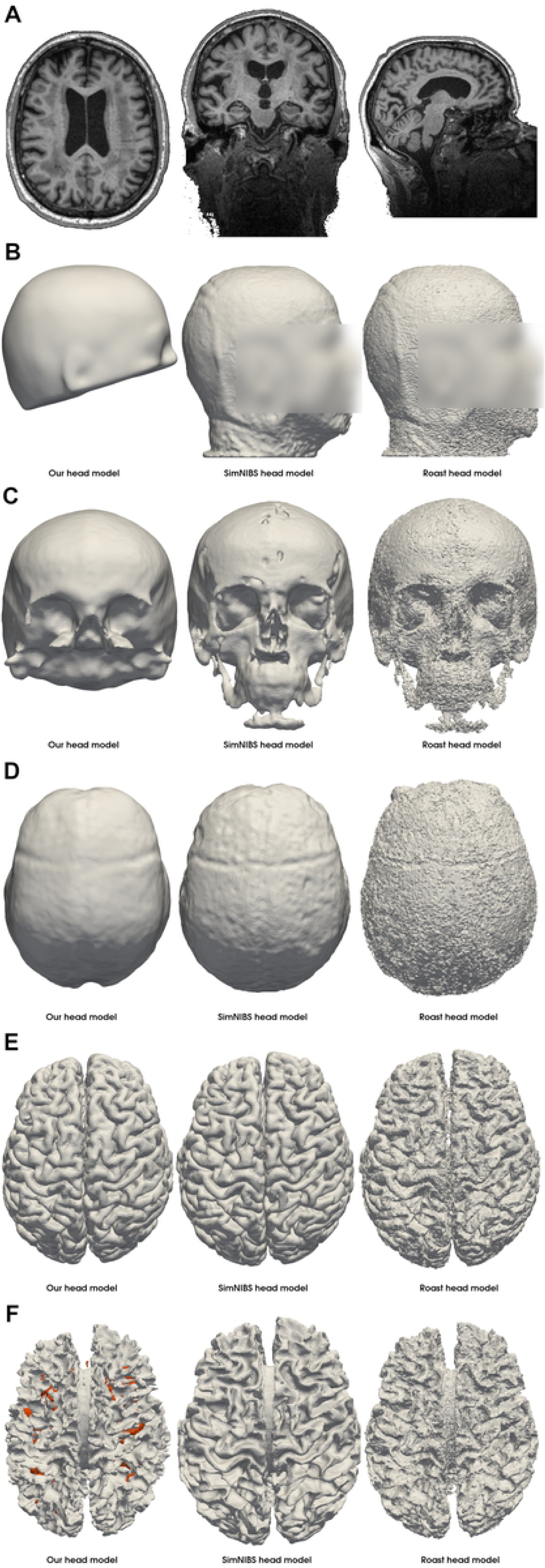
Mesh compartments of the head model generated using imaging data of the LIFE study (47). The T1-weighted imaging data **(A)** of a subject from the local, large-scale cross-sectional imaging study, LIFE, were used to create the head model using our approach, SimNIBS 3.0 and ROAST 2.7.1. Our approach yielded the most robust skull segmentation **(C)**. The skin compartment **(B)** is highly smoothed while maintaining the basic shape. The cerebrospinal fluid **(D)** and gray matter **(E)** mesh compartments are comparable across all three approaches. We included white matter lesions (**F**, orange), which were segmented from an additional T2-FLAIR image, into the white matter compartment of our head model.

**Fig 19.**
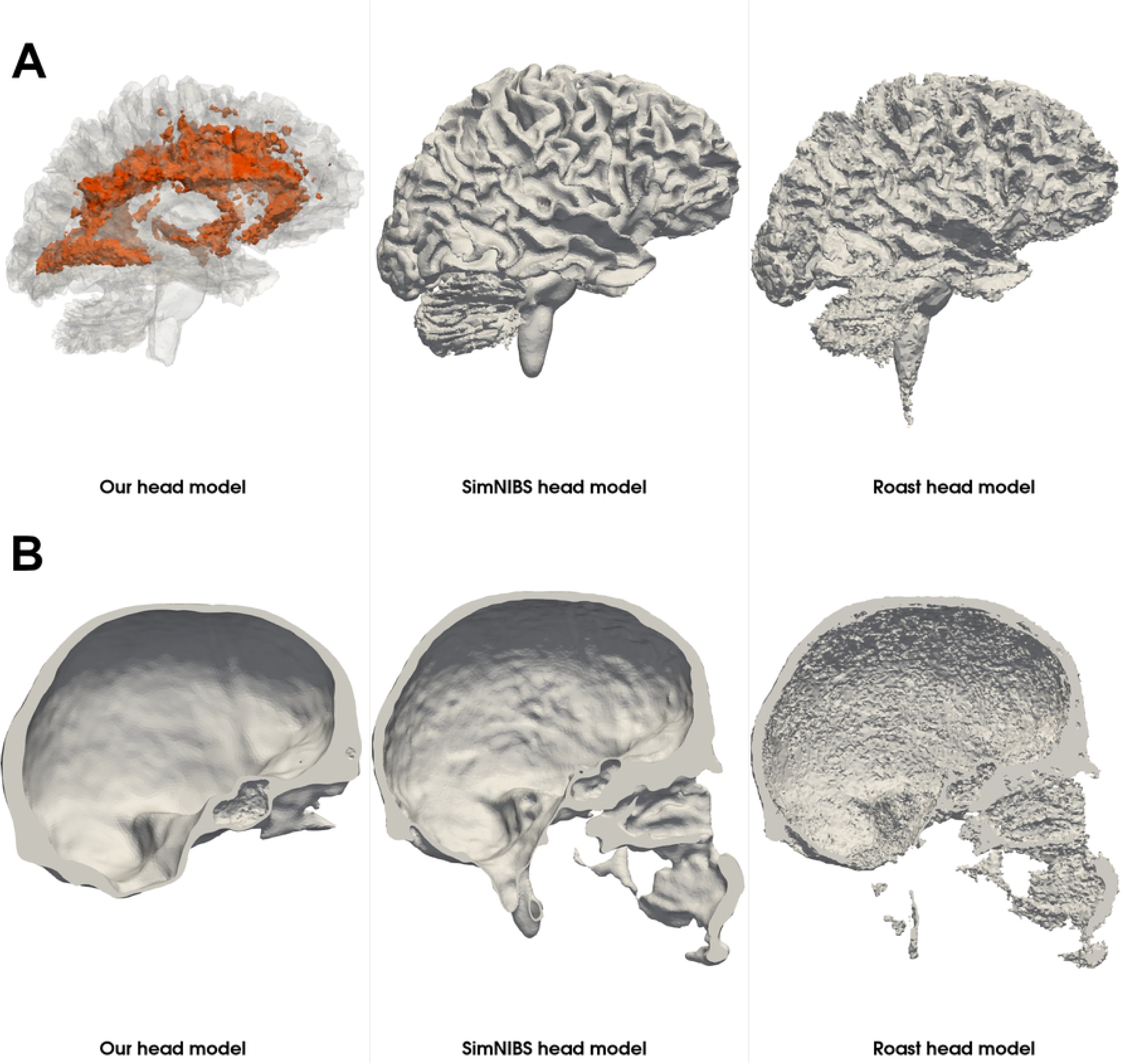
Visualization of the white matter lesions and skull thickness. The white matter lesions of a subject exhibiting a high lesion load are highlighted in orange **(A)**. Our atlas-based approach for skull segmentation tends to overestimate the thickness of the skull occipitally, along the superior sagittal sinus **(B)**.

We conducted a tDCS simulation using the generated white matter lesion head model with the following parameters: a bihemispheric setup of quadratic 5 cm by 5 cm electrodes as before, a 2 mA input current strength, default conductivity values from Table 2 for the standard tissue and 0.05 S/m for the lesioned tissue, modeling a calcification of the tissue. We then simulated the test case again assigning the conductivity of healthy white matter to the lesioned tissue. Comparing both computed electrical fields reveals a local perturbation in the area of the lesions (Fig. 20).

**Fig 20.**
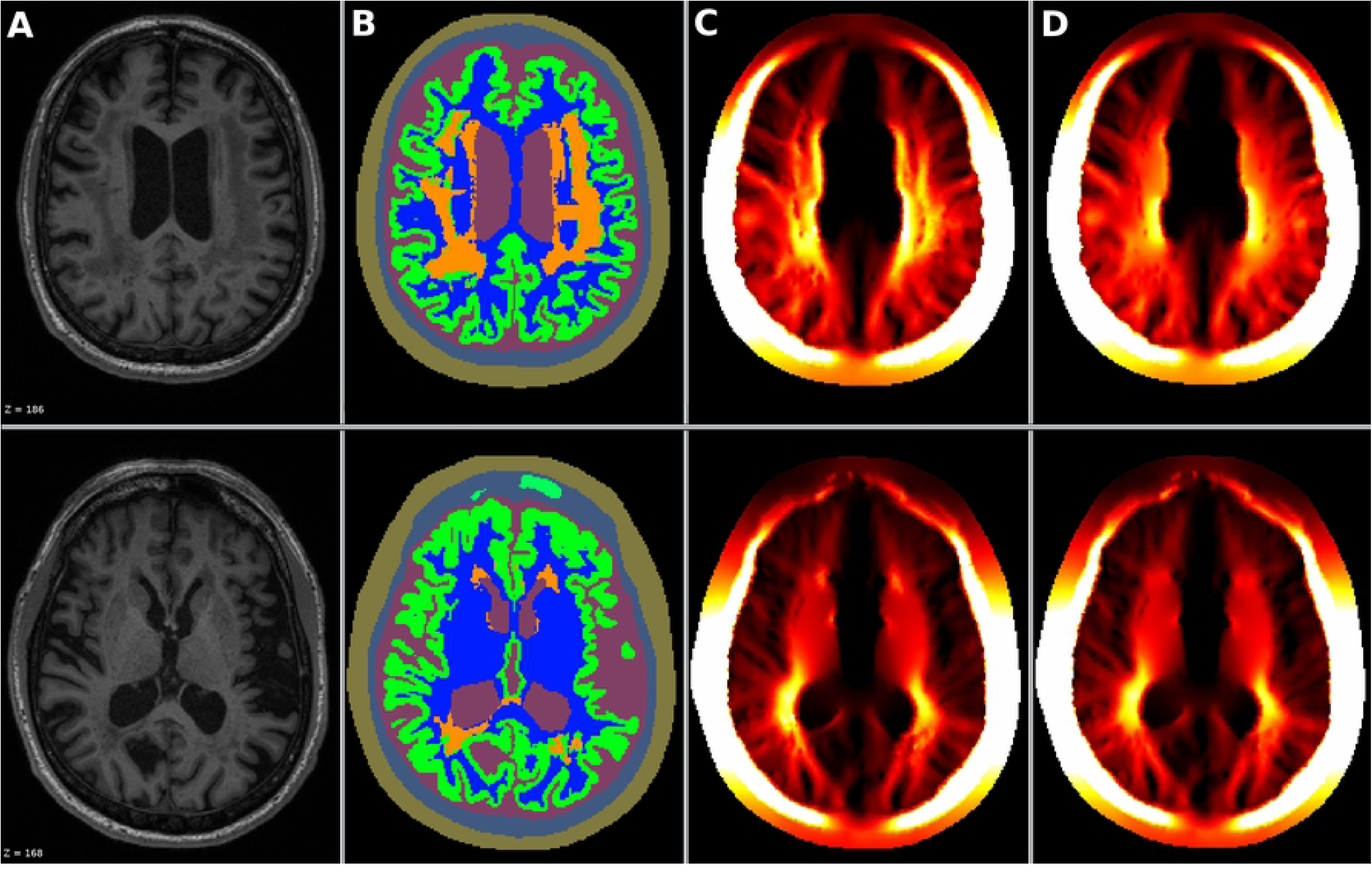
Simulation results using a white matter lesion head model. From the individual MR-image of subject **(A)** with a high lesion load (Fazekas score: 3) we segmented the standard tissue (skin, skull, csf, gray matter, white matter, air) and the white matter lesions **(B)**. We simulated **(C)** a bihemispheric electrode setup with quadratic 5 cm x 5 cm electrodes and a low conductivity of 0.05 S/m for the lesioned tissue while all other tissues were set to their default values (Table 2). For comparison we simulated again with the conductivity of the lesioned tissue set to that of healthy white matter **(D)**. A local perturbation of the electrical field in the area of the lesions can be observed.

## 4 Discussion

We presented a set of approaches for an individualized simulation of transcranial electric stimulation. The entire workflow from segmentation, meshing, electrode modeling, simulation, and visualization is built around OpenFOAM, a finite-volume based framework for numerical simulations. A coupled use, as well as the use of single features, are equally possible. Essential features are: 1) Individual head models are created solely from T1-weighted MRI data. Despite the limited T1-contrast we robustly segment scalp, skull, subarachnoid CSF, the ventricles, GM, WM, and the air cavities in the skull, as demonstrated using an exemplarily head image from a local, large-scale imaging study (47). 2) Combining image-based meshing with the surface-based meshing preserves the feature edges of the electrodes while avoiding any restrictions concerning the topology of tissue structures of the head model. 3) Arbitrary electrode shapes can be modeled, and their positioning is standardized according to the international 10-20 system. 4) Anisotropic tissue conductivity can be incorporated into the simulation. We demonstrated an overall agreement with an analytical 3-layer sphere model and the simulation results obtained by the simulation pipeline SimNIBS, especially when the simulations are based on the same head model, allowing comparability of the simulation results across simulation studies.

The combination of an image-based and a surface-based meshing algorithm realizes the head model generation. The image-based meshing holds two advantages. First, there is no restriction concerning the topology of the sub-compartments of the mesh. As the boundaries are determined directly from a labeled image, there is no requirement of overlap-free boundaries of sub-compartments (50). Therefore, the inclusion of structures that do not obey a strictly nested arrangement, for example, tumorous or lesioned tissue or holes in the skull (12), is facilitated. Second, image-based meshing is less sensitive to the quality of the input data which avoids extensive postprocessing of the segmentation images. However, boundaries may be less accurately approximated which we mitigated by setting a strict tolerance of the involved bisection algorithm. Surface-based meshing approximates boundaries most accurately and can preserve feature edges, which is, therefore, beneficial for representing any structure that does not require the flexibility of the image-based meshing, especially for the electrodes. As a consequence of the combination of both approaches, a tetrahedral volume mesh of high quality with maximum flexibility concerning the topology and maximum geometrical accuracy is obtained.

Comparing the results of our solver application and the solver employed in SimNIBS using an identical head model indicated an overall agreement in the global distribution and changes of the electrical field strength. However, peak differences of up to 66.2 % and peak deviations in the local electrical field direction of up to 40.6° were revealed in sparse locations close to the electrodes while on average the differences with the gray matter mesh compartments remained relatively small (approximately 15% difference in the field magnitude and 11° in local field direction). Since the volume mesh, the boundary conditions, and the conductivity values for the individual mesh compartments were identical, we conclude that differences arose due to the fundamentally different numerical approaches used for solving Maxwell’s equation (SimNIBS: finite-element method, OpenFOAM: finite-volume method).

We consider the finite-volume method implemented in OpenFOAM more sensitive to the quality of the volume mesh than the finite-element method in SimNIBS. Only by applying a limited interpolation scheme for the Laplacian term of the underlying equation of the electrical potential, the solution converged slowly when solving the tES problem in the head models created by SimNIBS. This choice of the discretization scheme resulted in a decreased convergence and thereby an increased solution time of approximately 4 minutes as compared to 100 seconds using our mesh with an even higher number of tetrahedra (SimNIBS mesh ≈ 4 M., our head model ≈ 5 M.). In addition, we partly attribute the observed differences between the solutions of the FVM solver and the FEM solver to the chosen numerical scheme. Most notably our volume meshed contained in the worst-case approximately 220 non-orthogonal cells whereas the SimNIBS volume meshes exhibited more than 1000 non-orthogonal cells in the best case and had problematic cells with negative cell volume, high skew, wrong orientation and a high aspect ratio as detected by the checkMesh utility of OpenFOAM. The higher mesh quality is the result of an extensive mesh optimization phase in our meshing approach, which increases the time for the volume meshing to up to 3 hours as compared to 5 minutes without optimization.

Deviations in the electrical field strength when simulating with our version of the Almi5 and Ernie head models instead of the ready-to-use head model might originate from differences in the caudal extent of the head model (51) and a different segmentation of the white matter and especially of the skull (Fig. 9). While our approach for skull segmentation tends to overestimate the skull caudally and occipitally, along the superior sagittal sinus, it slightly underestimates the thickness dorsally where the electrodes are attached. The thinner skull in that region may yield an overall higher electrical field magnitude (50). However, the general agreement in the change of the magnitude of the electrical field strength indicates that our modeling workflow does not introduce unexpected alterations to the head model.

The Blender plugin provides powerful means for the positioning and the modeling of the electrodes. After manually defining four fiducial points (nasion, inion, tragi of the ears) electrodes are placed automatically according to the 10-20 system. Any position outside the 10-20 system can be manually defined by moving the electrode across the scalp surface. A standard rectangular electrode is automatically modeled at the specified position. Other electrode types such as ring electrodes or triangular electrodes as applied in (52) and (53) are respectively possible, but require an adaption of the automated workflow.

Our solver application was verified using an analytical 3-layered sphere model and by comparison of the simulation results with the established simulation pipeline SimNIBS. However, a verification of the obtained simulation results with in-vivo recordings of the electrical field remains an open task. Promising approaches are electrical current density measurements obtained by the means of magnetic resonance electrical impedance tomography (54) or in-vivo recordings of the electrical potential by intracranial electrodes. TDCS simulations have been validated using intracranial recordings of epilepsy patients before (55), (56).

Since our workflow mainly focuses on addressing individual problems that we faced during the simulation of tDCS, it only provides a loose framework for the coupling of the suggested tools. These tools may, therefore, be easily interchanged by other tools or extended by new functionalities. However, familiarization with the individual tools and knowledge about the information flow between the tools (Fig. 1) is necessary to apply the workflow as a whole and potentially results in a higher initial effort for the setup and application as compared to fully automatized pipelines (10) (12).

First simulation studies suggest that damaged brain tissue due to a stroke influences the field distribution (57). Considering pathological tissue in the head model is, therefore, a vital extension to apply tES simulations to stroke patients. The coupled tools and the image-based meshing of our approach are prepared for this application as we demonstrated by the inclusion of white matter lesions into the head model. However, a fully automated and reliable segmentation of these irregular structures, especially stroke lesions, is still an open task for future research.

The advantageous properties of our suggested approaches for head and electrode modeling, as well as segmentation, facilitate simulation studies investigating alternative electrode shapes or irregular structures of the head model such as lesions and tumors in patients, implants, holes in the skull or vascular tissue.

## Acknowledgments

Benjamin Kalloch was funded by FAZIT-STIFTUNG and the International Max Planck Research School on the Neuroscience of Communication (IMPRS NeuroCom).

